# p53 mRNA availability during the cellular stress response is controlled by Cdc73 sequestration to stress granules

**DOI:** 10.1101/2022.05.25.493446

**Authors:** Hojin Lee, Tae-Hyeon Kim, Joo-Yeon Yoo

**Affiliations:** Department of Life Sciences, Pohang University of Science and Technology (POSTECH), Cheongam-Ro 77, Pohang, Geyongbuk, 37673, South Korea

## Abstract

Cells trigger the assembly of stress granules (SGs) under various stress conditions. Among the many proteins recruited to SGs are RNA-binding proteins (RBPs) and regulators of transcription. Here, we report the translocation of hCdc73, a component of the PAF1 transcription complex, to cytosolic SGs in response to sodium arsenite, MG132, thapsigargin (TG) or heat treatment. The hCdc73 protein possesses a long intrinsically disordered region (IDR) from amino acids 256-416, the presence of which is required and essential for the translocation of hCdc73 to cytosolic SGs. The purified hCdc73 IDR formed droplets in vitro, and the light-activated assembly of hCdc73 IDR-Cry2 was also verified. Alone, the hCdc73 IDR, however, was not sufficient for the translocation of hCdc73 to SGs, as physical interactions with the scaffold proteins of SGs, such as FMR1, were needed. Selective sequestration of cytosolic hCdc73 into SGs coincided with the dissociation of p53 mRNA from the hCdc73/Ski8/eEF1Bγ complex, resulting in a transient rise in p53 mRNA at the posttranscriptional level. In conclusion, we propose that in addition to the storage of nontranslating mRNAs, SGs also function to control the availability of mRNAs for stress response genes by restraining their negative regulators within SGs.

## INTRODUCTION

Organisms have evolved ways to respond to internal and environmental stresses for the maintenance of homeostasis and survival (1,2). The process collectively known as the cellular stress response is triggered by various hazardous conditions, including the presence of oxidants, genotoxins, metabolic inhibitors, or extreme temperatures (3,4,5,6). In response to stress, cells activate a signaling cascade to suppress global levels of translation and to reorganize gene expression by selectively inducing the expression of a specific set of genes from a pool of presynthesized mRNAs (7,8). This is an efficient and safe way to respond immediately to cellular stress signals without a significant time delay in new mRNA synthesis.

Under cellular stress conditions, mRNAs that are not undergoing translation and stalled mRNAs preferentially bind various RNA-binding proteins (RBPs) and are stored within nuclear bodies or cytoplasmic bodies, the assembly of which is controlled by liquid–liquid phase separation (LLPS) (9,10,11). One of the most studied types of cytoplasmic bodies that function during the cellular stress response is the stress granule (SG). Nontranslating mRNAs and translation initiation components are found within SGs, suggesting the storage function of SGs during stress responses. SGs are expected to influence gene expression by directly modulating translation through the relocation of essential regulatory proteins for translation and mRNAs away from the protein synthesis machinery (12,13). The assembly of SGs is also directly induced by translational arrest, as the release of bulky polysome-free mRNAs triggers SG assembly along with RBPs, thereby protecting nontranslating mRNAs from degradation (14).

Recent transcriptomic and proteomic studies of SGs have confirmed that stalled translation factors, in addition to RNAs and various RBPs, are major constituents of SGs (15). In contrast to the projected role of SGs in global mRNA protection, however, relatively small levels of mRNAs are actually enriched within SGs (16,17). Furthermore, reporter mRNAs designed to localize to SGs are translated and degraded at rates similar to those of their cytosolic counterparts (18.19). Although stress-induced compartmentalization of ribonucleoprotein (RNP) granules provides an effective means of spatiotemporal regulation of gene expression, how RNP granules selectively orchestrate the expression of stress-responsive genes is not clear. Interestingly, regulators of transcription and nucleocytoplasmic transporters are also included among the proteome of SGs (20,21,22,23). As evidenced by the neurodegenerative behavior of nucleocytoplasmic transport factors sequestered in SGs (24), the selective recruitment of transcription regulators into SGs suggests that they might function to specifically control the expression of stress-responsive genes. It has also been reported transcription regulators with neurodegenerative mutations sequester themselves within SGs, which disrupts their nuclear localization and function (25).

Parafibromin/hCdc73 is one of the components of the human RNA polymerase II (RNAPII)-associated factor complex (PAFc). PAFc is a complex of Ctr9, Paf1, Leo1 and Rtf1, in addition to Cdc73. Ski8, a component of RNA exosomes, was also found to be associated with PAFc in humans (26,27,28). Through protein–protein interactions, PAFc recruits transcription regulatory proteins and chromatin modifiers to target genomic loci and is thereby involved in almost every step of transcription mediated by RNA Pol II (26,29). A recent cryo-EM study of PAFc illustrated a loosely interconnected conformation of PAFc in which each component interacts with a different subunit of RNA Pol II. While the Leo1-bound form of Paf1 and Ctr9 sit on the outer boundary of RNA Pol II, the association of Cdc73 with PAFc or RNA Pol II is less stable, and Cdc73 even seems mobile (30).

In support of this discovery, the PAFc-independent behavior of Cdc73 has been widely reported. hCdc73 acts as a nuclear platform/scaffold protein to integrate and convert signals conveyed by the Wnt, Hedgehog, or Notch morphogen pathways, depending on its phosphorylation status (31,32). hCdc73 physically associates with the mRNA processing complex consisting of cleavage and polyadenylation specificity factor (CPSF) and cleavage stimulation factor (CstF), which is necessary for proper 3’ end modification of mRNAs in the nucleus (33). hCdc73 also forms a stable complex with eukaryotic elongation factor 1B (eEF1Bγ) and hSKi8 in the cytosol (34). Through its interaction with Ski8 and eEF1Bγ, which directly binds the mature form of p53 mRNA, hCdc73 controls the stability of p53 mRNA and p53-mediated apoptosis (34). In addition to its canonical role in translation, the RNA-binding properties of eEF1Bγ allow it to play a pivotal role in transcription, especially in p53-mediated cellular stress responses (35).

hCdc73 has been observed within the nuclear amyloid body, the protein assembly of which is induced by cellular stress signals (36). Moreover, Cdc73 has also been identified among the proteomes of cytosolic SGs (20, 37). Here, we experimentally demonstrate that cytosolic hCdc73 translocates to SGs and that its translocation contributes to the spatiotemporal regulation of p53 mRNA under cellular stress conditions.

## RESULTS

### hCdc73 translocates to SGs under various cellular stress conditions

In a previous study, we observed that hCdc73 in the cytosol functions to regulate the stability of mature p53 mRNA via its interaction with eEF1Bγ and Ski8 (34). Since hCdc73 is normally located and functions in the nucleus as a component of the PAF complex, we hypothesized that the cytosolic localization and related function of hCdc73 in the cytosol might be subject to control. To search for physiological conditions that affect the cytosolic localization of hCdc73, we tested various stress signals that induce the formation of SGs. Upon the treatment of cells with sodium arsenite, a representative reagent that causes oxidative stress and the unfolded protein response (UPR), or MG132, an inhibitor of 26S proteasomal degradation, a slight increase in the amount of cytosolic hCdc73 was detected (Figure 1A). At the same time, the transient assembly of cytosolic hCdc73 was observed under fluorescence microscopy, and cytosolic hCdc73 coincided with the typical marker proteins of SGs, G3BP1 and TIAR (Figure 1B, Supplementary Figure 1A). In addition to arsenic stress, we tested the effect of heat shock stress and ER stress by incubating cells at 46°C for 1 h or by treating cells with thapsigargin (TG), respectively (Figure 1A, B, Supplementary Figure 1B). Although the number and morphology of SGs differed, we consistently observed the colocalization of cytosolic hCdc73 where G3BP1 or TIAR assembled (Figure 1B). The translocation of cytosolic hCdc73 to SGs at various concentrations of sodium arsenite was separately verified (Supplementary Figure 1C). In addition to HeLa cells, we confirmed the localization of hCdc73 within SGs in HEK-293 and U-2OS cells (Supplementary Figure 1D).

**Figure 1.**
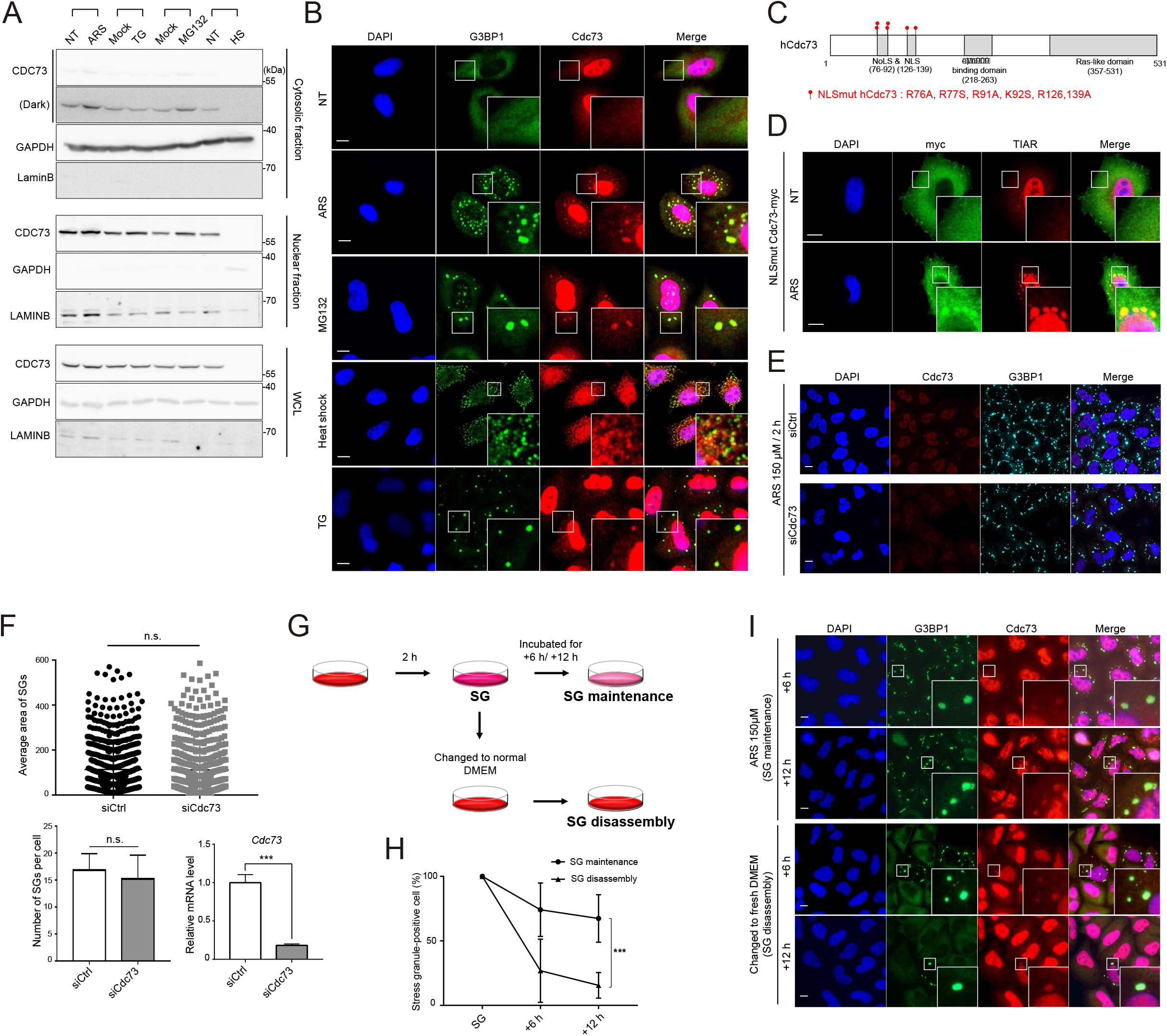
hCdc73 translocates to stress granules (SGs) under various stress conditions. (A) Immunoblot analysis of hCdc73 from the cytosol (top), nucleus (middle), and whole-cell lysates (bottom). HeLa cells were treated with 500 μM sodium arsenite for 1 h (ARS), 50 μM thapsigargin for 1 h (TG), 10μM MG132 for 4 h (MG132), and 46°C heat shock for 1 h (HS). Each sample was loaded next to its own control set; NT: nontreated, Mock: DMSO-treated. GAPDH and LAMINB were used as marker proteins for the cytosol and nuclear fractions, respectively. (B) Confocal images of endogenous hCdc73 and G3BP1 in cells left untreated (NT) or treated, as indicated. The same conditions in (A) were used. (C) Schematic diagram of the mutation sites used to generate NLSmut hCdc73. (D) Confocal images of NLSmut hCdc73-myc-transfected HeLa cells; untreated (NT) or treated with 500 μM sodium arsenite for 1 h (ARS). TIAR, marker for SGs. (E) Confocal images of endogenous hCdc73 and G3BP1 in siCdc73- or siCtrl-transfected HeLa cells treated with 150 μM sodium arsenite for 2 h. (F) Area (Top) and number of SGs per cell (Bottom left) quantitated from the data in (E) (n=51). (Bottom right) Knockdown efficiency of siCdc73 was measured by RT-qPCR. (G) Experimental schematic describing the SG maintenance and disassembly conditions. (H) Percentage of SG-positive cells measured during SG maintenance and disassembly (n=50-99). (I) Representative confocal images of endogenous hCdc73 and G3BP1 tested at +6 h and +12 h of SG maintenance and disassembly. All scale bars in this figure, 10 μm. Scatter plot graph and bar graphs were presented as mean ± standard deviations. Statistical values were calculated using an unpaired t-test.

The localization of cytosolic hCdc73 within SGs was also tested with exogenously transfected hCdc73-myc. Although the colocalization of hCdc73-myc where G3BP1 assembles was observed, the overall intensity of cytosolic hCdc73-myc was weak (Supplementary Figure 1E). It seems that expression of the transfected hCdc73-myc was overrepresented in the nucleus. To increase the amount of cytosolic hCdc73 protein, we generated mutant forms of hCdc73 in which its nuclear and nucleolar localization sequences were mutated (Figure 1C). hCdc73 possesses a functional nuclear localization signal (NLS) from amino acids 126-139 and a nucleolar localization signal (NoLS) from amino acids 76-92 (38,39). As expected, the mutation of both NLS and NoLS (NLSmut) resulted in the preferential expression of hCdc73 in the cytosol (Figure 1D). hCdc73 (NLSmut)-myc-expressing cells were then stimulated with sodium arsenite, and the colocalization of hCdc73 (NLSmut)-myc proteins where the SG marker protein TIAR assembled was observed, similar to the colocalization of endogenous hCdc73. Observing the colocalization patterns of hCdc73 and SGs, we wondered whether hCdc73 plays any role in the formation of SGs. However, when the expression of hCdc73 was silenced, the number of G3BP1-positive SGs was unaffected, indicating that hCdc73 is unlikely to function during the SG assembly process (Figure 1E, F). Instead, it seems that hCdc73 is recruited to SGs when they assemble.

The formation of SGs is a reversibly regulated process, as the assembly and disassembly of SGs are dynamically controlled. We therefore questioned what would happen to assembled hCdc73 when SGs disassembled. To address this question, HeLa cells were incubated with 150 μM sodium arsenite for 2 h and then cultured in normal culture medium for an additional 6 or 12 h to remove stress inducers (Figure 1G, “SG disassembly”) or were continuously maintained in sodium arsenite-containing medium (Figure 1G, “SG maintenance”). During recovery periods in normal medium, the percentage of SG-containing cells dramatically declined, indicating that SGs had disassembled. After 12 h of recovery in normal medium, only 14% of cells possessed SGs (Figure 1H, Supplementary Figure 1F). In contrast, when cells were continuously maintained in sodium arsenite-containing medium, as a control, sustained assembly of SGs (64%) was observed, even after 12 h of incubation (Figure 1H, Supplementary Figure 1F). Using these experimental systems, we examined the localization of endogenous hCdc73 relative to SGs. In accordance with the disappearance of SGs, the punctuate form of cytosolic hCdc73 also disappeared. As a result, assembled cytosolic hCdc73 was detected only when SGs were present (Figure 1I). These data indicate that hCdc73 not only translocates to SGs but also follows the dynamics of SGs in the cytosol.

### An intrinsically disordered region (IDR) within the hCdc73 protein is required for its translocation to SGs

To understand how hCdc73 translocates to or assembles within SGs, we attempted to identify the domains responsible for hCdc73 that control this behavior. hCdc73 contains a C-terminal Ras-like domain that shares homology with yCdc73, and known interaction domains for β-catenin or for eEF1Bγ /Ski8 lie within the N-terminal region (34,40,41). In addition, that most cancer-related mutations are observed within the N-terminal region of hCdc73 indicates the functional importance of this region (42). Although the crystal structures of the C-terminal domain of yCdc73 and the N-terminal half (amino acids 1-111) of hCdc73 have been separately reported (40,43), the crystal structure of the full-length form of hCdc73 is not available, suggesting that unstructured low-complexity regions might be present within hCdc73. To address this hypothesis, we utilized the D^2^P^2^ program (44) to search for low-complexity regions within hCdc73 (Supplementary Figure 2A). Broad regions encompass amino acids 256-416 scored high for low complexity; of these residues, the region could be further subdivided into ‘IDR(N)’, which covers the first N terminal half of the IDR (amino acids 256-360), and the ‘IDR(C)’, covering the second IDR domain (amino acids 361-416) (Figure 2A). As the assembly of SGs depends on the LLPS properties of intrinsically disordered proteins, we focused on these putative IDR domains of hCdc73. Thus, hCdc73 mutants that lack the individual IDR(N) or IDR(C) region, or both, were generated and tested for their capability to translocate into SGs (Figure 2A, B). Unlike the full-length form of hCdc73, all of the hCdc73 deletion mutants tested failed to translocate to SGs. To quantitate the colocalization of hCdc73 with SGs, we calculated the SG enrichment score, which measures the fluorescence intensity of hCdc73 within SGs (Figure 2C). Next, we examined whether the IDR alone was sufficient for translocation to SGs. Thus, truncated forms of hCdc73 containing IDR(N), IDR(C), or both were generated and tested (Figure 2A, D). Under arsenic stress conditions, the translocation of IDR(N+C) and IDR(N) of hCdc73 to SGs was observed, as clear colocalization with TIA-1, a marker protein of SGs, was detected. However, IDR(C) of hCdc73 behaved differently. Overexpression of IDR(C) of hCdc73 alone was sufficient to form puncta in the absence of arsenic stress (Figure 2D). Furthermore, IDR(C) of hCdc73 did not colocalize with SGs when SG formation was induced (Figure 2D, E). These data suggest that the IDR of hCdc73 has dual functions. While IDR(N) of hCdc73 is essential for the translocation of hCdc73 into SGs, the effect of the IDR(C) region is somewhat complicated, as this region was needed but not sufficient to exert translocation into SGs upon arsenic stress.

**Figure 2.**
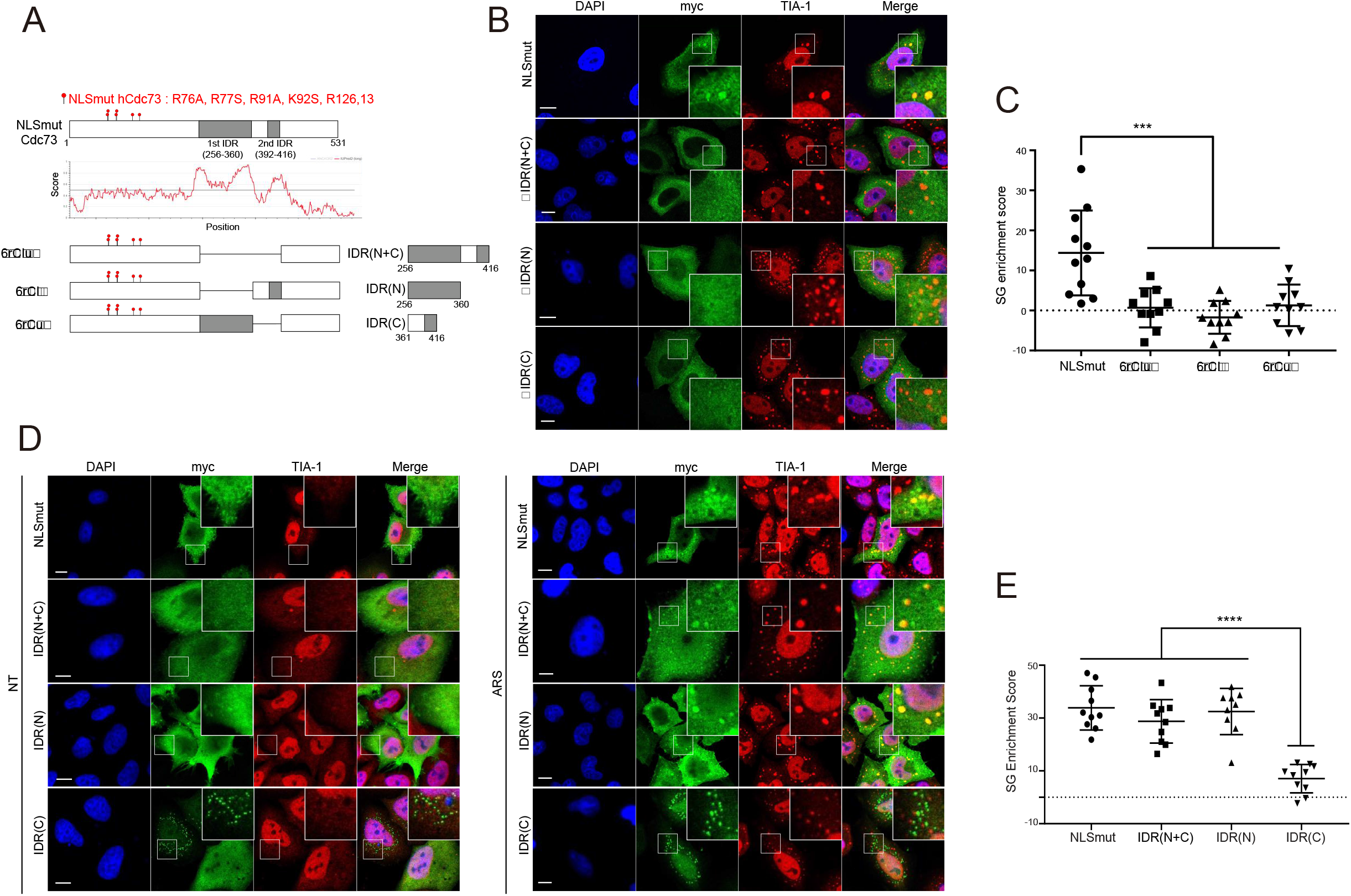
The intrinsically disordered region (IDR) within the hCdc73 protein is required for its translocation to SGs. (A) Schematic diagram of various hCdc73 IDR mutants. Presented IDR score of hCdc73 obtained from the IUPred simulation program (60). The disordered region is indicated by a gray box. (B, D) Representative confocal images of transfected hCdc73 IDR mutants. Left, untreated (NT) or treated with 500 μM sodium arsenite for 1 h (ARS). TIA-I, marker for SGs. (C, E) Quantification of hCdc73 IDR mutants within SGs (n=9-11). The SG enrichment score of each hCdc73 IDR mutant was measured as described in the ‘Materials and methods’ section. All scale bars in this figure, 10 μm. Scatter plot graphs were presented as mean ± standard deviations. Statistical values were calculated using a Turkey’s multiple comparisons test.

### The phase separation properties of hCdc73-IDR are required but not sufficient for the translocation of hCdc73 to SGs

Proteins with intrinsically disordered properties can undergo phase separation to form membraneless organelles. In addition to the scaffolding protein of SGs, G3BP1/2, many proteins that assemble within SGs tend to possess IDRs and form liquid droplets (15,45,46). Therefore, we tested whether the IDR of hCdc73 also possesses the ability to undergo phase separation and whether this property is needed for the translocation of hCdc73 to SGs. To investigate the phase separation properties of hCdc73-IDR, recombinant 6x His-GFP-tagged IDR(N+C) of hCdc73 was purified and tested for its ability to form droplets *in vitro*. With the aid of a crowding reagent (10% PEG 8000), the purified protein at 10 μM formed liquid droplets under physiological buffer conditions (150 mM NaCl, pH 7.4) (Figure 3A). The assembly of liquid droplets was sensitive to the salt concentration, as liquid droplets were barely detected under high-salt (300 mM NaCl) conditions. Acidic or basic buffer conditions also similarly prevented droplet assembly (Figure 3B). Under these pH conditions, hCdc73-IDR(N+C) seemed to aggregate. When the purified hCdc73-IDR(N+C) protein was treated with 1,6-hexanediol at pH 7, the droplets disappeared, indicating that weak hydrophobic interactions are important for this assembly event (Figure 3A). Of note, the aggregates observed at pH 5 and pH 9 were resistant to 1,6-hexanediol treatment.

**Figure 3.**
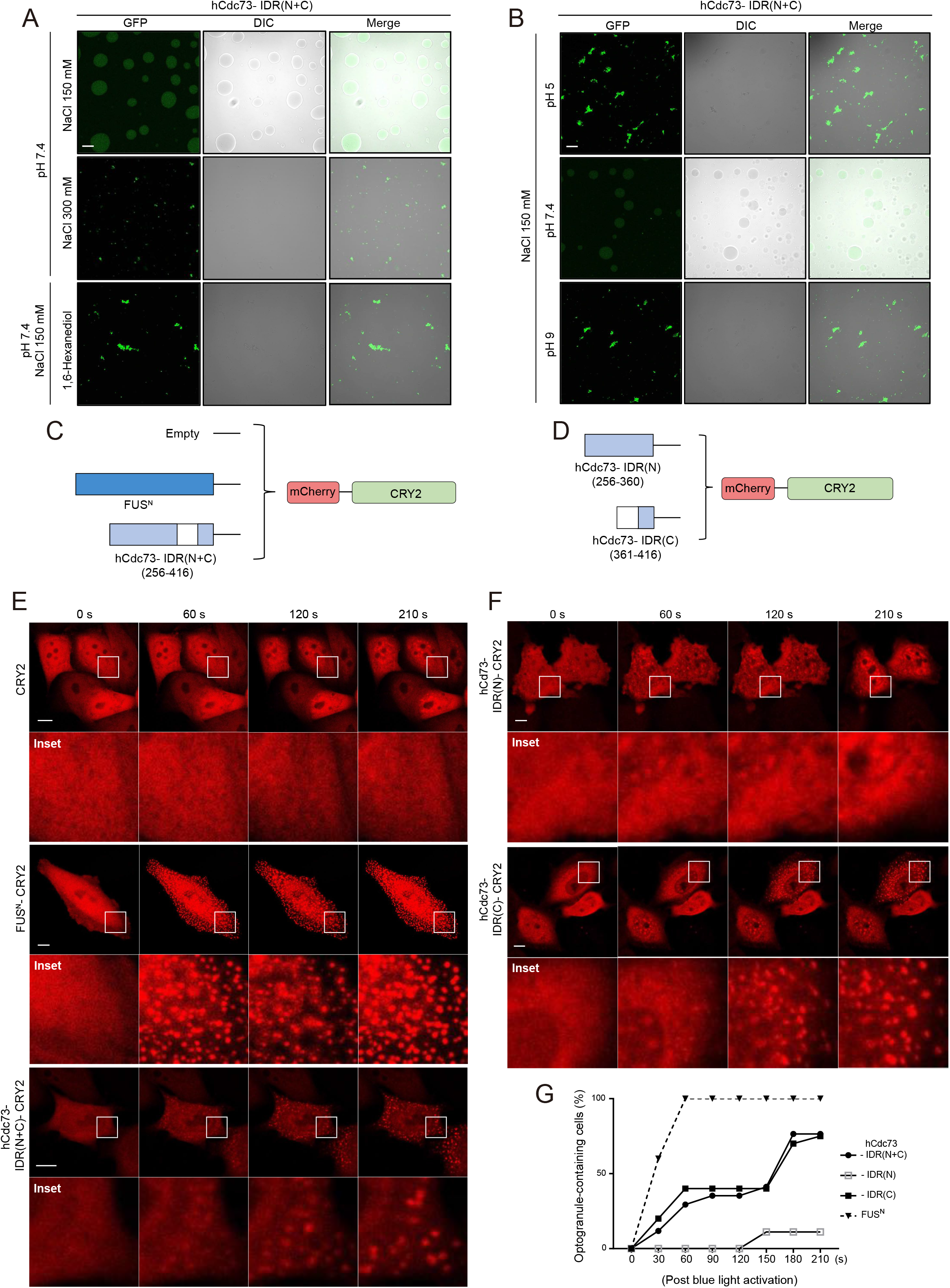
Phase separation properties of hCdc73 IDR. (A, B) Representative images of the *in vitro* droplet formation assay. Recombinant IDR(N+C)-GFP (10 μM) was incubated at the indicated pH and salt concentration with 10% PEG8000. 1,6-Hexanediol (5%) was applied, where indicated, to disturb weak hydrophobic interactions. Scale bar, 10 μm. (C, D) Schematic diagram of the constructed optoDroplet plasmids. As a positive control, the IDR of FUS (FUS^N^) was used. (E, F) Time-lapse imaging of droplet formation upon blue light exposure in HeLa cells. Scale bar, 10 μm. (G) Quantification of optoDroplet-containing cells; data from (E) and (F) (n=5-20).

Separately, we also tested the droplet-forming ability of hCDC73-IDR in a cell system using optoDroplet techniques (47). For this, mCherry-hCdc73 IDR(N+C) was fused to Cry2, a light-sensitive self-associating protein, and the light-inducible phase separation properties of hCdc73-IDR were examined after blue light exposure (Figure 3C). As a positive control, the disordered N-terminal domain of FUS (FUS^N^; amino acids 1-214) that covers the prion-like domain was also tested. Upon blue light (488-nm laser)-induced activation, mCherry-hCdc73 IDR(N+C)-Cry2 underwent light-inducible phase separation, resulting in distinct droplet assembly (Figure 3E). In contrast to FUS^N^, however, the kinetics of droplet assembly were rather slow. Under our experimental conditions, the phase separation of mCherry-FUS^N^-Cry2 was detected after 30 sec of exposure, and most cells formed droplets after 60 sec of light exposure (Figure 3E, G). However, light-induced droplets of mCherry-hCdc73 IDR(N+C)-Cry2 were barely detected after 30 sec of exposure and started to appear after 60 sec of exposure. Even after longer light exposure, not every cell tested formed droplets (Figure 3E, G), indicating that hCdc73 IDR(N+C) is able to undergo phase separation but possesses relatively weak activity.

Next, we tested whether hCdc73 IDR(N) and hCdc73 IDR(C) possess intrinsic phase separation properties utilizing the optoDroplet system (Figure 3D, F). While mCherry-hCdc73 IDR(C)-Cry2 exhibited phase separation properties similar to those of mCherry-hCdc73 IDR(N+C)-Cry2, mCherry-hCdc73 IDR(N)-Cry2 was not able to form droplets (Figure 3F, G). These data suggest that the phase separation properties of hCdc73 IDR depend on IDR(C). Without IDR(C), hCdc73 did not colocalize with SGs (Figure 2B, C), indicating that the phase separation properties of hCdc73 IDR are required for its translocation. However, hCdc73 IDR(C) alone failed to translocate to SGs (Figure 2D, E), again indicating that the phase separation properties of hCdc73 IDR are not sufficient for its translocation to SGs. Earlier, we observed that the presence of IDR(N) was both required and sufficient for the recruitment of hCdc73 to arsenic stress-induced SGs (Figure 2C, E). Based on these data, we hypothesized that IDR(N) of hCdc73 possesses other essential properties for SG translocation, such as specific protein–protein interactions required for recruitment.

### The carrier protein FMR1 is required for the translocation of hCdc73 to SGs

SGs are composed of ‘core’ or ‘scaffold’ proteins surrounded by less concentrated ‘shell’ or ‘client” proteins, along with various mRNAs (20,48). Since the phase separation properties of hCdc73 IDRs are necessary but not sufficient for the recruitment of hCdc73 within SGs, we speculated that hCdc73 likely behaves as a client rather than as a core scaffolding protein. In general, the recruitment of client proteins to SGs requires specific protein–protein interactions with scaffolding proteins (20). Therefore, we first tested whether hCdc73 physically interacts with G3BP1, a major core protein of SGs. However, we did not observe a physical interaction between these proteins (Supplementary Figure 3A, B). Therefore, we next asked whether the recruitment of hCdc73 to SGs requires physical interactions with carrier proteins, which are heavily associated with SGs. In our search for putative carrier proteins associated with SGs, we first selected 6 candidate proteins that are abundant and frequently studied (49). HeLa cells were separately transfected with siRNAs targeting each candidate protein, and the colocalization of cytosolic hCdc73 to sodium arsenite-induced SGs was examined (Figure 4A-C). Among the tested proteins, the silencing of FMR1 showed the most dramatic effect, as the recruitment of cytosolic hCdc73 to sites where G3BP1 assembles was significantly decreased (Figure 4C). Although gene expression was repressed accordingly, the silencing of CAPRIN1, FXR1, NUFIP2, UBAP2L, or USP10 did not dramatically change the colocalization patterns of hCdc73 and SGs. These data suggest that FMR1 functions to aid, probably via physical interactions, hCdc73 for its location within SGs. This hypothesis was also strengthened by a recent paper describing the proteomic analysis of FMR1 (50). To address whether FMR1 indeed physically interacts with hCdc73, we performed a GST pull-down assay using the lysates of Flag-FMR1- and GST-hCdc73-transfected HeLa cells. hCdc73 interacted with the full-length form of the FMR1 protein (Figure 4D). Moreover, hCdc73 IDR(N) was sufficient for a physical interaction with FMR1. These data led us to conclude that hCdc73 interacts with FMR1 through the IDR(N) domain. Since we previously observed that IDR(N) of hCdc73 is essential for the translocation of hCdc73 into SGs (Figure 2C, E), we concluded that FMR1, as a carrier protein, brings hCdc73 to SGs via physical interaction.

**Figure 4.**
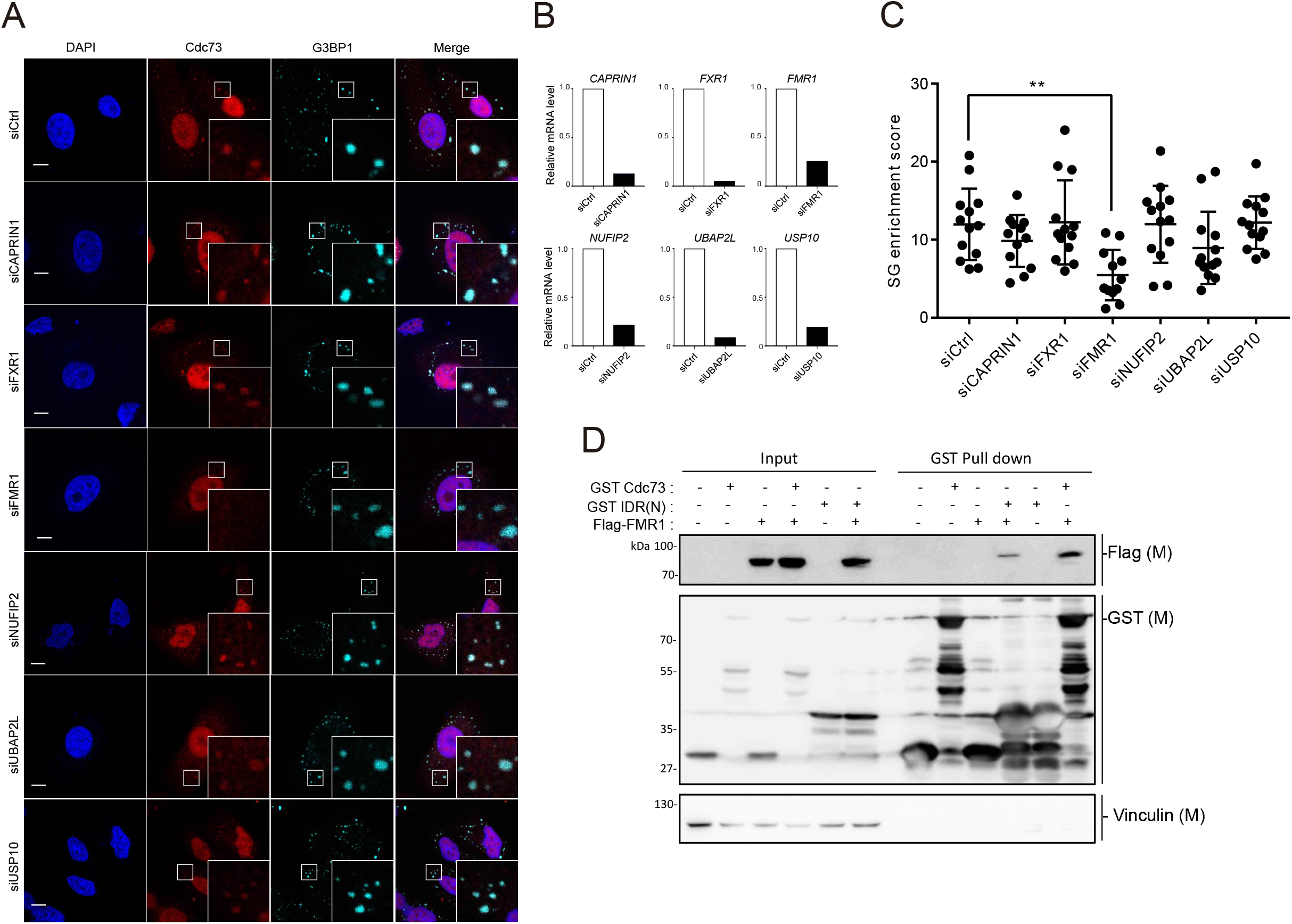
The translocation of hCdc73 to SGs requires the scaffold protein FMR1. (A) Representative confocal images of endogenous hCdc73 and G3BP1 in siRNA-transfected HeLa cells as indicated treated with 150 μM sodium arsenite for 2 h. Scale bars, 10 μm. (B) Knockdown efficiency measured by RT–qPCR. (C) SG enrichment scores obtained from the data in (A) (n=12-13). Scatter plot graph was presented as mean ± standard deviations. Statistical values were calculated using a Dunnett’s multiple comparison test (D) HEK-293T cells were transfected with the indicated plasmids and subjected to a GST pulldown assay.

### p53 mRNA levels and the formation of SGs are positively correlated

SG-inducing cellular stressors trigger global translational arrest that results in a high concentration of nontranslating RNAs in the cytosol (12,13). The compartmentalization of RNAs within SGs is predicted to provide a protective shelter for ribosome-free RNAs. However, a recent study also suggested that SG formation is an adaptive response that selectively renders stress-responsive mRNAs available (51). Since we previously observed that cytosolic hCdc73 controls the stability of p53 mRNA, we wondered whether the abundance of p53 mRNA changes during the process of SG assembly and whether hCdc73 plays a role in this regulation. To test this idea, we first examined whether p53 mRNA levels are altered under stress conditions that induce the assembly of SGs. HeLa cells were treated with various stress inducers, and p53 mRNA levels were measured. To focus on immediate changes in mRNA levels, instead of transcriptional controls, most of the stress inducers were applied acutely. Under various stress conditions, we observed statistically significant changes in p53 mRNA levels, except under TG-induced stress conditions (Figure 5A). Similarly, acute treatment with sodium arsenite and MG132 increased p53 mRNA levels up to 1.5-fold compared to those in mock-treated controls. Acute heat shock stress also slightly increased p53 mRNA levels, but the effect was marginal. The effect of sodium arsenite at different doses on p53 mRNA levels was also verified in HeLa cells or in HEK-293 and U-2OS cells (Supplementary Figure 4A, B).

**Figure 5.**
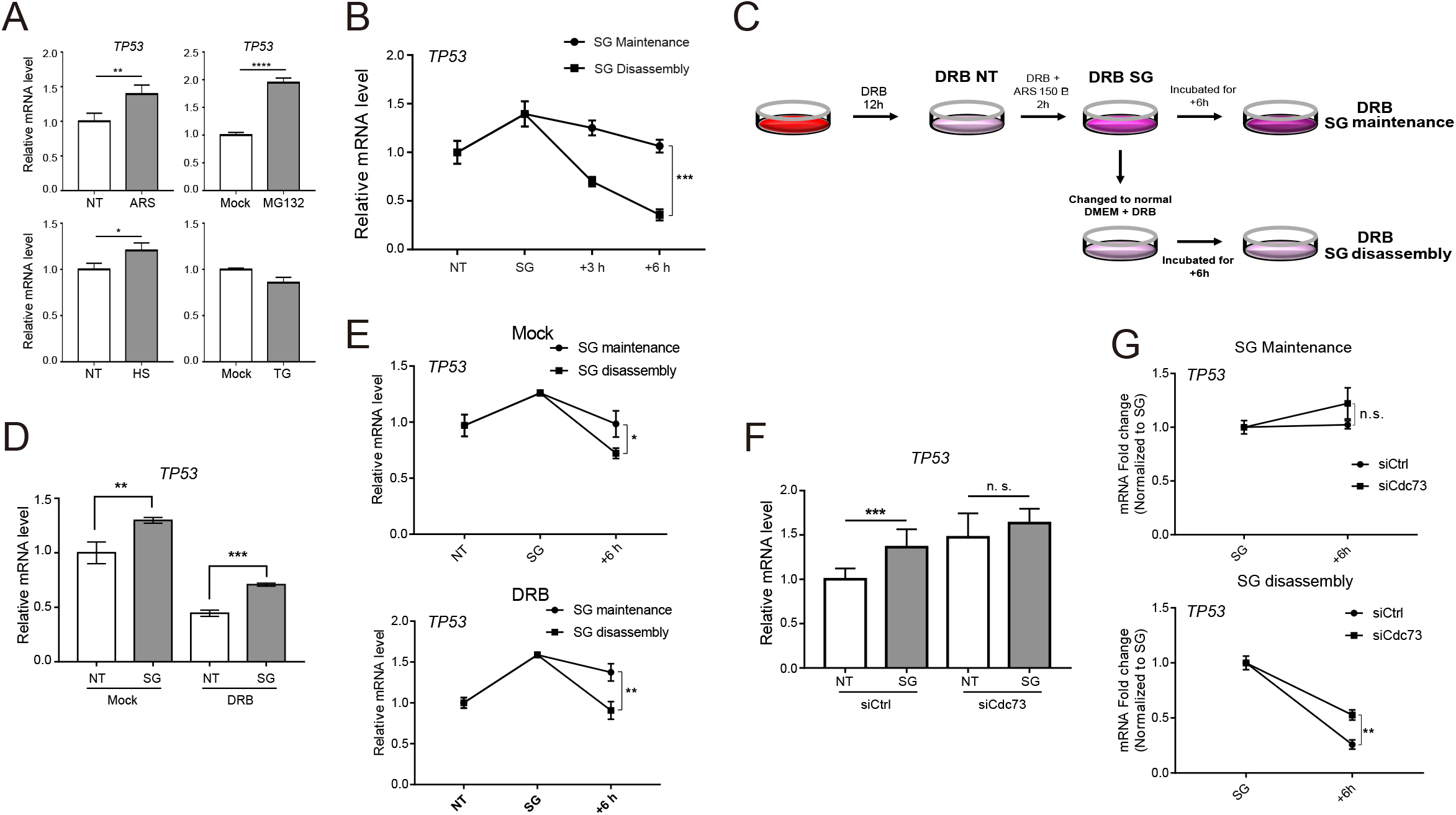
p53 mRNA levels and SG assembly are positively correlated. (A) HeLa cells were treated with 500 μM sodium arsenite for 1 h (ARS), 10 μM MG132 for 4 h, 50 μM thapsigargin for 1 h (TG), and 46°C heat shock for 1 h (HS). The p53 mRNA level in each stressed cell type was measured by RT–qPCR and compared with its own control set (NT: nontreated, Mock: DMSO-treated). (B) HeLa cells were treated with 250 μM sodium arsenite for 1 h (SG), followed by SG maintenance or disassembly. At the indicated times, p53 mRNA levels were measured by RT–qPCR. (C) Experimental protocol with DRB. After 12 h of DRB pretreatment, the cells were acutely stimulated with 150 μM sodium arsenite for 2 h, followed by SG maintenance or disassembly, including DRB treatment. (D-E) HEK-293 cells were treated with DRB and sodium arsenite as indicated in (C). (F-G) Control or hCdc73 siRNA-transfected HeLa cells were acutely treated with sodium arsenite (150 μM, 2 h), followed by SG maintenance or disassembly. p53 mRNA levels were measured by RT–qPCR. Each graph represents mean data from three independent experiments and error bars are presented as the mean ± standard deviations. Statistical values were calculated using an unpaired t-test.

To address whether the formation of SGs and changes in p53 mRNA levels correlate, we next examined changes in p53 mRNA levels during SG maintenance or disassembly. When cells remained under sodium arsenite-induced stress conditions for an additional 3 and 6 h, the increased p53 mRNA level was relatively sustained (Figure 5B, Supplementary Figure 4C), in accordance with the proportions of SG-forming cells (Figure 1H). In contrast, when arsenic-stressed cells were exposed to normal medium for the disassembly of SGs, p53 mRNA levels dramatically declined (Figure 5B, Supplementary Figure 4C).

Although changes in mRNA levels during acute (1 h) treatment are more likely due to posttranscriptional regulation, we could not exclude the possibility of transcriptional control. Therefore, cells were pretreated with ‘5,6-dichlorobenzimidazole 1-β-D-ribofuranoside (DRB), an inhibitor of transcription, and the formation of SGs and levels of p53 mRNA were examined. We chose to pretreat cells with 100 μM DRB for 12 h since it halved the level of p53 mRNA (Figure 5C, Supplementary Figure 4D) but did not alter sodium arsenite-induced SG formation (Supplementary Figure 4E). Upon arsenic treatment, p53 mRNA levels rose to similar levels in the absence or presence of DRB (Figure 5D). Additionally, we measured the amounts of p53 mRNA upon culture under SG maintenance and disassembly conditions with DRB pretreatment. The data obtained from the DRB-treated cells were similar to those for the mock-treated control cells (Figure 5E). These data together indicate that changes in p53 mRNA levels under sodium arsenite-induced stress conditions are regulated at the posttranscriptional level.

Finally, we tested the involvement of hCdc73 in the regulation of p53 mRNA levels under arsenic stress conditions. To address this question, p53 mRNA levels were examined in arsenic-stressed, hCdc73-silenced cells. As reported, hCdc73 silencing alone resulted in enhanced p53 mRNA levels (34). Upon arsenic stress, however, further induction of p53 mRNA was not detected in hCdc73-deficient cells (Figure 5F), suggesting that arsenic stress might counterbalance the negative effect of hCdc73 on p53 mRNA. Separately, the effect of hCdc73 on the p53 mRNA level was also tested under SG maintenance and disassembly conditions (Figure 5G). Under SG maintenance conditions, the changes in p53 mRNA levels were not significant, regardless of whether hCdc73 was depleted. On the other hand, the decline in the p53 mRNA level during SG disassembly was substantially delayed in the hCdc73-depleted cells, indicating that the degradation of p53 mRNA during the recovery phases requires the presence of hCdc73.

### hCdc73, but not 53 mRNA or eEF1Bγ, translocates to arsenic stress-induced SGs

Our data thus far demonstrated that 1) p53 mRNA levels increase at the posttranscriptional level when SGs assemble, 2) hCdc73 is required for this mRNA regulation, and 3) hCdc73 is sequestered to SGs. Since the cytosolic hCdc73/eEF1Bγ/Ski8 complex associates with p53 mRNA for degradation control (34), these observations led us to hypothesize that the entrapment of hCdc73 within SGs might work to prevent p53 mRNA degradation and to increase the cytosolic pool of p53 mRNA. To understand how this regulation occurs, we examined what happened to the hCdc73/eEF1Bγ/Ski8 complex and p53 mRNA during SG assembly. To do so, we first performed a fluorescence in situ hybridization (FISH) assay to visualize the spatial distribution patterns of p53 mRNA under arsenic stress conditions. A p53-specific RNA probe detected a broad p53 mRNA distribution in untreated cells. Upon sodium arsenite treatment, the spatial distribution pattern of p53 mRNA was not changed, and moreover, significant colocalization with G3BP1-positive SGs was not detected (Figure 6A, Supplementary Figure 5A). Among the mRNAs reported to translocate to SGs (16,17), we selected CDK6 mRNA as a positive control for FISH experiments. As expected, CDK6 mRNA colocalized well with G3BP1-positive SGs upon sodium arsenite treatment (Figure 6B, Supplementary Figure 5B). Therefore, these data suggest that p53 mRNA is not included among the many other mRNAs stored within SGs during stress. A search for SG transcriptome data also revealed that p53 mRNA was not enriched in SGs (16,17).

**Figure 6.**
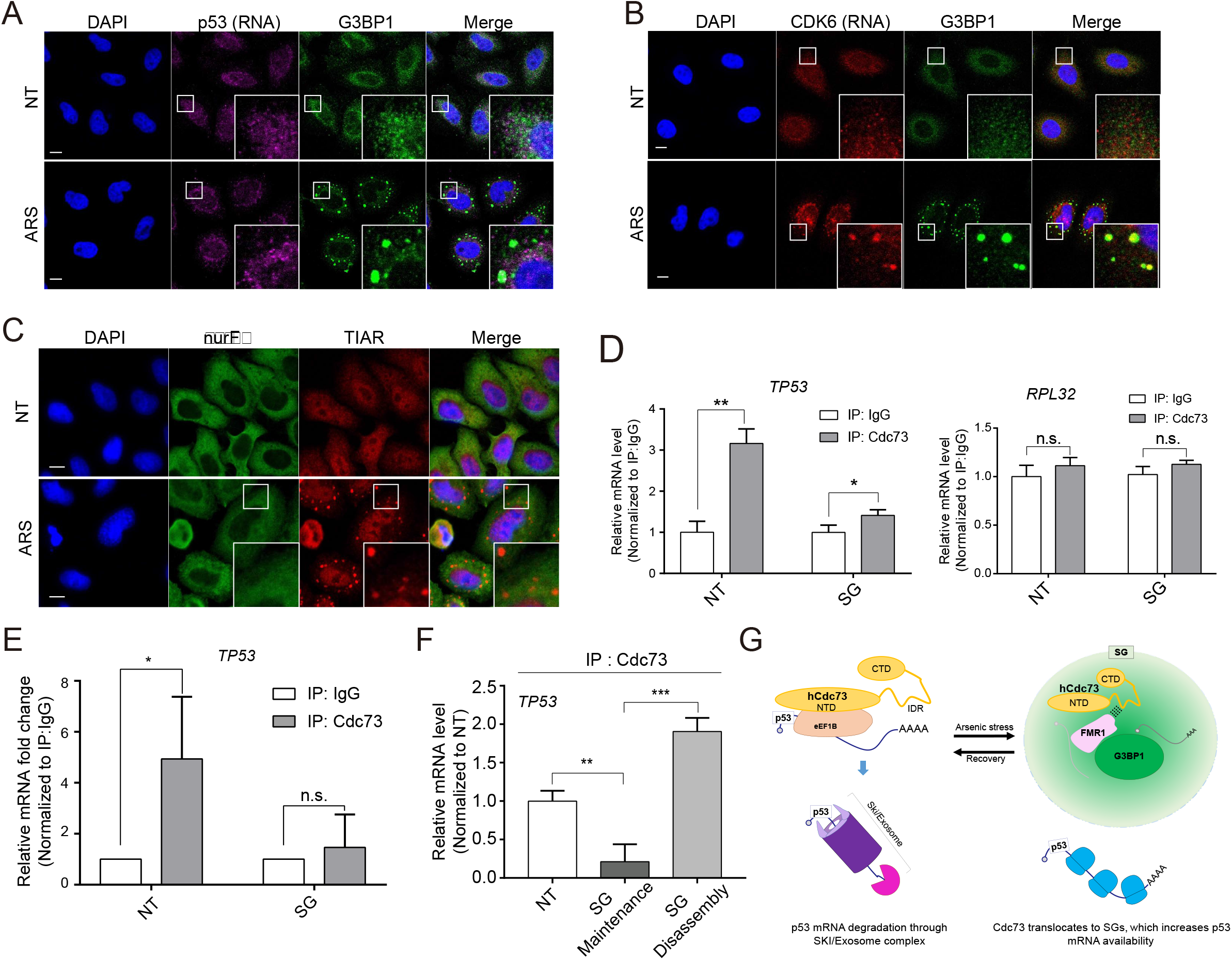
Neither p53 mRNA nor eEF1Bγ translocates to SGs. (A-B) Confocal images from RNA fluorescence in situ hybridization (RNA FISH) experiments. HeLa cells were either untreated (NT) or treated with 500 μM sodium arsenite for 1 h. FISH probes that specifically bind p53 (A) and Cdk6 (B) mRNAs were used. G3BP1 was used as a marker for SGs. Scale bars, 10 μm. (C) Confocal images of endogenous eEF1Bγ or TIAR in HeLa cells. Left untreated (NT) or treated with 500 μM sodium arsenite for 1 h (ARS). Scale bars, 10 μm. (D) RNA-IP was performed using IgG or endogenous hCdc73 antibodies and the cytosolic fraction of HEK-293 cells. Left untreated (NT) or treated with 500 μM sodium arsenite for 1 h (ARS). Bound mRNA was measured by RT–qPCR; p53 (left) or hRPL32 (right). (E) RNA-IP was performed using IgG or endogenous hCdc73 antibodies and the cytosolic fraction of HeLa cells (crosslinked with 0.03% formaldehyde). Left untreated (NT) or treated with 500 μM sodium arsenite for 1 h (ARS). Cytoplasmic hCdc73-bound p53 mRNA was measured by RT– qPCR, and the fold change was measured. (F) Results of RNA-IP using endogenous hCdc73 antibody. HeLa cells were treated with 500 μM sodium arsenite for 1 h and then incubated for 6 h under SG maintenance or disassembly conditions (NT: untreated). (G) Graphical abstract of hCdc73- and SG-mediated p53 mRNA control. Each graph represents mean data from three independent experiments and error bars are presented as the mean ± standard deviations. Statistical values were calculated using an unpaired t-test

Next, we examined spatial information of the eEF1Bγ protein during arsenic stress. eEF1Bγ was of particular interest, as the association of hCdc73 with p53 mRNA depends on this protein (34). Unlike hCdc73, however, the cellular localization pattern of the eEF1Bγ protein did not change dramatically upon sodium arsenite-induced SG formation, and colocalization with SGs was not detected (Figure 6C). Therefore, these data indicate that hCdc73 dissociates from the eEF1Bγ-p53 mRNA complex and translocates to SGs under arsenic stress conditions. To confirm this hypothesis, we performed an RNA-immunoprecipitation (RNA-IP) assay to detect cytosolic hCdc73 and p53 mRNA using the cytoplasmic fraction (Figure 6D). The data clearly showed that the interaction between hCdc73 and p53 mRNA was significantly decreased under sodium arsenite-induced SG assembly conditions. When tested in SG disassembly conditions, the interaction between hCdc73 and p53 mRNA increased, again confirming that the entrapment of hCdc73 within SGs works to prevent the association of hCdc73 with p53 mRNA (Figure 6E,F). Based on these data, we concluded that the selective sequestration of hCdc73 away from the cytoplasmic RNP complex containing p53 mRNA functions to transiently increase the availability of p53 mRNA under cellular stress conditions (Figure 6G).

## DISCUSSION

The tumor suppressor p53 is a transcription factor (TF) involved in the cellular decision of death and survival. It functions to inhibit cell proliferation and promote cell death when the cell is faced with detrimental cellular stress. Additionally, the activity of p53 is required for the maintenance of metabolic pathways that contribute to cell survival (52). For this reason, the abundance and activity of the p53 protein are tightly controlled during diverse stress conditions (53). In addition to p53 production, the stability of p53 mRNA and protein are subject to control for more acute responses. Previously, we reported that hCdc73 forms a novel complex with eEF1Bγ and Ski8 that functions to degrade p53 mRNA in the cytosol (34). In this report, we demonstrate a novel way to posttranscriptionally control p53 mRNA levels under cellular stress conditions by the dissociation of hCdc73 from the eEF1Bγ/Ski8-p53 mRNA cytosolic complex and its sequestration into SGs (Figure 6G).

We show that p53 mRNA levels positively correlate with SG levels. Under stress conditions in which SGs assemble, enhanced p53 mRNA levels were observed (Figure 5A). In accordance with SG disassembly, decreased p53 mRNA levels were detected (Figure 5B). These changes in p53 mRNA levels during SG formation were independent of transcriptional activity, similar to the changes in p53 mRNA levels detected when DRB, an inhibitor of transcription, was applied (Figure 5D, E). Additionally, we demonstrated that, unlike many other mRNAs that are stored within SGs during stress, p53 mRNA remained in the cytosol (Figure 6A). Under these conditions, the physical association of hCdc73 with p53 mRNA was disrupted (Figure 6D). Based on these observations, we proposed that the selective sequestration of hCdc73 into SGs is critical for enhancing p53 mRNA levels under cellular stress conditions, as its presence along with eEF1Bγ/Ski8 mRNA in the cytosol leads to p53 mRNA degradation.

Our data suggest that the translocation of hCdc73 into SGs is a critical step for the control of p53 mRNA availability under cellular stress conditions. For hCdc73 to be incorporated into SGs, the intrinsically disordered properties of hCdc73 as well as physical interactions with carrier proteins such as FMR1 are needed. Our investigation of the IDR domain of hCdc73 and its ability to form liquid droplets implies that the function of hCdc73 can be coordinated through concentration into biomolecular condensates. SGs are composed of many essential scaffold proteins, ‘client’ proteins recruited by physical interactions with scaffolding proteins or carrier proteins, and various kinds of RNAs (20). As hCdc73 is intrinsically disordered and localized within SGs, we initially wondered whether hCdc73 plays a role in the assembly of SGs. However, we did not observe a change in SG assembly when hCdc73 was silenced (Figure 1F). Furthermore, carrier proteins such as FMR1 are needed for the recruitment of hCdc73 within SGs (Figure 4A, B). Based on these observations, we postulate that hCdc73 is simply translocated to SGs in response to cellular stress. It seems that stressed cells induce the translocation of cytoplasmic hCdc73 to SGs to regulate p53 mRNA availability for a proper stress response.

From yeast to mammals, it is essential to reprioritize genes for transcription, from housekeeping genes to stress-responsive genes, upon exposure to harmful environmental conditions. Similar to p53 mRNA control by the selective sequestration of hCdc73 within SGs during cellular stress conditions, Pab1-mediated heat shock-responsive gene regulation depends on the phase separation properties of Pab1. In yeast, the mRNAs of heat shock-responsive genes are bound by Pab1, which inhibits the translation of heat shock-responsive genes under normal cellular conditions. Under conditions of heat shock stress, Pab1 forms a biomolecular condensate, which frees the mRNAs of heat shock-responsive genes for translation (51). Unlike Pab1, which directly binds mRNA, the physical association of hCdc73 with p53 mRNA requires the RBP eEF1Bγ. However, the cytosolic distribution pattern of eEF1Bγ did not change upon SG formation, and eEF1Bγ remained in the cytosol along with p53 mRNA (Fig 6C). Although the detailed mechanisms differ, it seems that a mechanistically conserved stress response controls the availability of mRNA via selective sequestration of mRNA-associated cytosolic proteins into phase-separating granules.

Our data collectively show that p53 mRNA levels are posttranscriptionally regulated during cellular stress. Although p53 is a master regulator of cellular stress responses, relatively little is known about its regulation by SGs. When a transcriptome analysis of SGs utilizing U-2OS cells stably expressing G3BP1-GFP was carried out, p53 mRNA was not enriched (16). In an analysis of the mRNAs that interact with the RBP Tia1, the dissociation of p53 mRNA in response to a DNA damage reagent was reported in activated B cells (54). Notably, increased USP10 in prostate cancer cells inhibits the p53 signaling pathway via interaction with G3BP2 (55).

Although our work mainly focused on cytosolic hCdc73, many studies on hCdc73 address its role as a component of the PAF complex in the nucleus. The PAF complex associates with RNAPII and functions as a protein scaffold via interactions with various TFs and transcriptional regulators (26,29). In the nucleus, spatiotemporal regulation of enhanceosomes that are undergoing LLPS functions for the dynamic recruitment of a transcriptional regulatory network (56,57). Coactivators and other regulatory factors of transcription form biomolecular condensates, the compartmentalization of which is driven by dynamic low-specificity interactions between IDRs and DNAs within the complex (56,58). In addition, the progression of RNAPII from the initiation stage to the elongation stage depends on its phase separation properties, which are controlled by hyperphosphorylation of the intrinsically disordered C-terminal domain (59). Discovery of the phase separation properties of hCdc73 via its IDR therefore opens the possibility of the involvement of hCdc73 or the PAF complex within nuclear biomolecular condensates with transcriptional activities. We also postulate that hCdc73 can be found in various molecular condensates in the nucleus, in addition to cytosolic SGs. Notably, nuclear hCdc73 was reported to be present within amyloid bodies, physiological amyloid aggregates that form within the nuclei of stressed cells (36).

In summary, here, we report the translocation of hCdc73 to SGs through the IDR domain of hCdc73 under various cellular stress conditions. The sequestration of hCdc73 also represents a novel way to control the mRNA availability of the stress-responsive p53 gene. Although we do not know how p53 mRNA is selectively targeted by cytosolic hCdc73, we are interested in determining how this selectivity is achieved and the repertoire of mRNAs that are controlled by this mechanism. As the activities of both nuclear and cytosolic hCdc73 are closely linked to carcinogenesis, it is also necessary to understand the physiological consequences of hCdc73 translocation to SGs and the relationship with tumorigenesis.

## MATERIALS AND METHODS

### Cell culture and transfection

HeLa, HEK-293, HEK-293T and U-2OS (ATCC, Manassas, Virginia) cells were grown in Dulbecco’s modified Eagle’s medium (DMEM; Lonza, Basel, Switzerland) supplemented with 10% fetal bovine serum (FBS; Welgene, Taipei, Taiwan) and routinely checked for mycoplasma contamination with a mycoplasma PCR detection kit (iNTRON, Seongnam, Korea). Lipofectamine 2000 (Invitrogen, Waltham, Massachusetts) and polyethyleneimine (PEI; Polysciences, Warrington, Pennsylvania) were used for transient transfection of short interfering RNA (siRNA) and plasmid DNA, respectively, according to the manufacturer’s recommendations. After 48 h of transfection with plasmid DNA, cells were harvested or fixed for further analysis. At 72 h post-transfection, cells transfected with siRNAs were harvested or fixed for further analysis.

### Plasmids, siRNAs, and oligomers

Myc-hCdc73 and GST-hCdc73 have been described previously (34). To make mutant forms of Myc-hCdc73, site-directed mutagenesis and Gibson assembly (NEB, Ipswich, Massachusetts) were performed. optoDroplet plasmids (a gift from Yongdae Shin, Department of Chemistry, Seoul National University, Seoul, Korea) were subcloned into pCDNA.3.1 for transient overexpression. To generate GST-IDR(N), IDR(N) was subcloned from Myc-IDR(N) into the pEBG vector. To generate flag-FMR1 and GST-eEFG1Bγ, HEK-293 complementary DNA was PCR amplified and cloned into the pFLAG-CMV and pEBG vectors, respectively. The siRNA and RT-qPCR primer sequences used in this study are provided in (Supplementary Table 1).

### Immunoblotting

For Western blot analysis, cells were lysed in lysis buffer (150 mM NaCl, 1% Triton X-100, 0.1% SDS, and 0.5% deoxycholic acid with protease inhibitors, 25 mM Tris-HCl, pH 7.5), and proteins in the total cell lysate were quantified using the Bradford method and separated on SDS-polyacrylamide gels at various percentages (7-12%). The proteins were transferred to nitrocellulose membranes and analyzed with the indicated antibodies. Finally, the membranes were incubated with horseradish peroxidase (HRP)-conjugated secondary antibodies, and an LAS4000 luminescent image analyzer (Fujifilm, Tokyo, Japan) was used to visualize immunoreactive signals. The signals were acquired using Supersignal West Femto Maximum Sensitivity substrate (Thermo Fisher Scientific, Waltham, Massachusetts).

### Cellular fractionation

For cytoplasmic and nuclear fractionation, 5×10^6^ cells were resuspended in sucrose buffer (320 mM sucrose, 3 mM CaCl_2_, 2 mM MgOAc, 0.1 mM EDTA, 0.5% Nonidet P-40, 1 mM dithiothreitol and 0.5 mM phenylmethyl sulfonyl fluoride, 10 mM Tris–HCl, pH 8.0). After centrifugation at 600 ×g, the cytoplasmic (supernatant) fraction was removed. The pellet was washed with sucrose buffer without Nonidet P-40 and lysed in lysis buffer. The mixture was incubated on ice for 30 min and centrifuged at 13,000 ×g. The supernatant, which contained the soluble nuclear proteins, was removed. The pellet was boiled with Laemmli sample buffer (Bio–Rad, Hercules, California) supplemented with 5% β-mercaptoethanol at 95°C for 10 min to obtain the insoluble fraction.

### Immunofluorescence microscopy and quantification

Cells were cultured on round cover glasses in 6-well culture dishes and washed with PBS, followed by fixation with cold methanol or 4% paraformaldehyde. The cells were then permeabilized by incubation with 0.2% Triton X-100 for 15 min and blocked with 5% BSA in PBST. The cells were incubated with primary antibodies overnight at 4°C, and then secondary antibodies conjugated to either Alexa Fluor 488, 568, or 647 (Invitrogen) in 1% BSA-PBST were incubated with the cells for one hour at room temperature. Then, a DAPI solution (Invitrogen) was used to visualize nuclei. Cover glasses were mounted on glass slides using fluorescence mounting medium (Agilent, Santa Clara, California) and analyzed with a FLUOVIEW FV3000 confocal microscope (Olympus, Tokyo, Japan) and TCS SP5 confocal microscope (Leica, Wetzlar, Germany). The area of SGs and the number of SGs were determined using the ImageJ plugin. The SG enrichment score was measured to quantitate the colocalization of hCdc73 and SGs. The area of SGs in the confocal microscopy images was set as the region of interest (ROI), and the fluorescence intensity of hCdc73 mutants within the ROI[SGs] was measured. Background ROI[BG] that exclude ROI[SGs] and represent the overall expression patterns of hCdc7 mutants in the cell were further designated. Next, the ‘SG enrichment score’ was calculated by dividing the intensity of hCdc73 mutants within ROI[SGs] to the intensity within the ROI[BG]. For imaging siRNA transfected cells, BLOCK-iT Fluorescent Oligo for Lipid Transfection (Invitrogen) was co-transfected with indicated siRNAs. Only fluorescent oligo-expressing cells are regarded as siRNA transfected cells.

### Purification of the IDR(N+C) of the hCdc73 protein

All purified proteins were expressed from the pET28a vector in BL21(DE3) *Escherichia coli* cells. Expression cultures were grown in LB broth with constant shaking (200-240 rpm) at 37°C for 10 h. The bacteria were incubated in culture medium at 18°C for 36 h to allow bacterial protein expression. Bacterial pellets were resuspended in binding buffer (20 mM Tris-HCl, 500 mM NaCl, 10% glycerol, 7 mM β-mercaptoethanol, 8 M urea, and 25 mM imidazole, pH 7.9) containing an EDTA-free protease inhibitor cocktail, followed by sonication to induce lysis. The supernatant obtained after centrifugation at 15,000 rpm for 30 min was purified using a HisTrap™ HP column (GE Healthcare, Chicago, Illinois). Next, the eluted proteins were dialyzed in dialysis tubing (Thermo Fisher Scientific) against a storage buffer (50 mM Tris-HCl, 0.5 M NaCl, 10% glycerol, and 7 mM β-mercaptoethanol, pH adjusted to 7.4). The proteins were further purified using size exclusion chromatography (Superdex 200 Increase 10/300 GL; GE Healthcare) to remove nonspecific proteins. Vivaspin® concentrators (Sartorius, Göttingen, Germany) were used to increase the concentration of the protein solution, which was subsequently stored at −80°C.

### In vitro phase separation assay

To assess LLPS, purified protein (10 μM) was mixed with LLPS buffer (150 mM NaCl, and 10% PEG8000, 50 mM Tris-HCl, pH 7.4). The salt concentration and pH were changed as indicated, and the effects of these parameters on IDR(N+C) droplet behavior were observed. Two microliters of protein mixed with LLPS buffer was loaded on 3% BSA-coated glass slides (Paul Marienfeld, Lauda-Königshofen, Germany) and covered with a 10ø cover slip (Deckglaser) to observe LLPS droplets. Images of the droplets were taken immediately after the proteins were mixed with LLPS buffer using an Axiovert 200M microscope (Zeiss, Oberkochen, Germany) or an FLUOVIEW FV3000 confocal microscope (Olympus). ZEN 3.0 or FV31S-DT software was used to analyze the images. To separate the supernatant and pellet according to the pH, purified protein was dialyzed in the pH buffer described below. Samples were centrifuged at 13,000 rpm for 1 min. The supernatant and pellet were separated and denatured in Laemmli buffer at 95°C for 10 min for SDS–PAGE, followed by Coomassie Blue staining or immunoblotting. Buffer pH 5 (20 mM sodium acetate, 150 mM, pH 5), buffer pH 7.4 (20 mM sodium phosphate, 150 mM NaCl, pH 7.4), buffer pH 9 (20 mM Tris-HCl, 150 mM NaCl, pH 9).

### optoDroplet assay

All images of optoDroplet plasmid-transfected HeLa cells were captured using a 40× objective and TCS SP5 confocal microscope (Leica) to observe IDR behaviour upon blue light activation (47). The intracellular distributions of the indicated IDRs were examined using a 561-nm laser following activation with blue light (488-nm laser). As a control, empty-mCherrry-CRY2-expressing cells were imaged with a 561-nm laser upon blue light activation (488-nm laser) for a maximum of 210 s. Afterward, optoDroplet-formed cells were quantitated.

### IP and GST pulldown assay

For IP, HEK-293T and HeLa cells were lysed with NP-40 lysis buffer (20 mM Tris-HCl at pH 7.4, 150 mM NaCl, 2 mM EDTA, and 1% Nonidet P-40) or polysome lysis buffer (PLB; 10 mM HEPES at pH 7, 100 mM KCl, 5 mM MgCl_2_, and 0.5% Nonidet P-40) (34). One milligram of cell lysate was incubated with mouse normal IgG or G3BP1 antibody (BD, Franklin Lakes, New Jersey, 611126) for 8 h with protein A/G agarose beads (Sigma-Aldrich, Burlington, Massachusetts) at 4°C. For the GST pulldown experiments, 1 mg of cell lysate was incubated with 40 μl of glutathione-Sepharose beads (GE Healthcare) at 4°C. The beads were then washed with lysis buffer five times, and proteins were eluted from the beads by incubation with Laemmli sample buffer supplemented with 5% β-mercaptoethanol at 95°C for 10 min. All the inputs were quantified using the Bradford method and loaded in equal amounts.

### RNA isolation and real-time reverse transcriptase–PCR

Total RNA was extracted using RNAiso Plus (Takara Bio, Shiga, Japan). RNA (1 μg) was reverse transcribed with ImProm-II Reverse Transcriptase (Promega, Madison, Wisconsin) using oligo(dT) or random primers (Promega). PCR analysis of the cDNA was performed on a StepOne Plus Real-Time PCR System (Applied Biosystems, Waltham, Massachusetts). In most of the PCR experiments, hRPL32 was used as the internal control. To inhibit transcription, 100 μM DRB (Sigma-Aldrich) was added for 12 h.

### FISH probes

The following FISH probes were purchased from LGC Biosearch Technologies (Lystrup, Denmark); TP53 (Stellaris® FISH Probes, Human TP53 with Quasar® 670 Dye) and CDK6 (Stellaris® FISH Probes, Human CDK6 with Quasar® 570 Dye). FISH was performed according to the manufacturer’s recommendations (Stellaris® RNA FISH Protocol for Adherent Cells).

### RNA-IP assay

Cells were either crosslinked with 0.03% formaldehyde (HeLa) or not crosslinked (HEK-293); afterward, RNA-IP was performed as previously described (34). Approximately 5×107 cells were lysed in PLB (100 mM KCl, 5 mM MgCl_2_, 10 mM HEPES, pH 7.0, and 0.5% Nonidet P-40) supplemented with protease inhibitors and 100 Uml-1 RNase inhibitor. After centrifugation at 3,000 ×g, the supernatant was precleared and then incubated with anti-hCdc73 antibody (Bethyl, Montgomery, Texas, a300-170a) and protein A/G agarose beads for 4 h at 4°C. After five washes with 1 ml of PLB, the bound complexes were eluted, and RNA was isolated from the eluate using RNAiso Plus.

### Antibodies and reagents

Reagents: sodium arsenite, Thapsigargin, MG132, and 1,6-hexanediol are purchased from Sigma-Aldrich.

Antibodies: hCdc73 antibody (Bethyl, a300-170a) was used for immunoblotting and IP. For immunofluorescence microscopy of hCdc73, antibodies from Invitrogen (PA5-26189) and Santa Cruz (Dallas, Texas, sc-33638) were used. SG marker antibodies against G3BP1 (BD, 611126), TIA-1 (Abcam, Cambridge, UK, ab40693), and TIAR (Cell Signaling Technology, Danvers, Massachusetts, 5137) were purchased as indicated. Primary antibodies against the following were used: GAPDH (Santa Cruz, sc-47724), LAMINB (Santa Cruz, sc-374015), C-MYC (Santa Cruz, sc-40), GST (Santa Cruz, sc-138), Vinculin (Santa Cruz, sc-55465), and Flag (Sigma-Aldrich, F7425).

### Statistical analysis

All statistical analysis were performed and graphed using GraphPad Prsim (version 9.2.0). The significance of differences was evaluated by an unpaired two-tailed t test or 1way ANOVA multiple comparisons. All data are presented as the mean ± standard deviations. Three independent experiments were conducted to perform statistics, unless otherwise noted in the corresponding figure legend. Significance is defined as *, p < 0.05; **, p < 0.005; ***, p < 0.0005; ****, p<0.00005; n.s., not significant.

## ACKNOWLEDGEMENT

This work was supported by the National Research Foundation of Korea(NRF) grant funded by the Korea government(MSIT)(NRF-2020R1A2C2009707)

## FIGURE LEGENDS

**Supplementary Figure S1.**
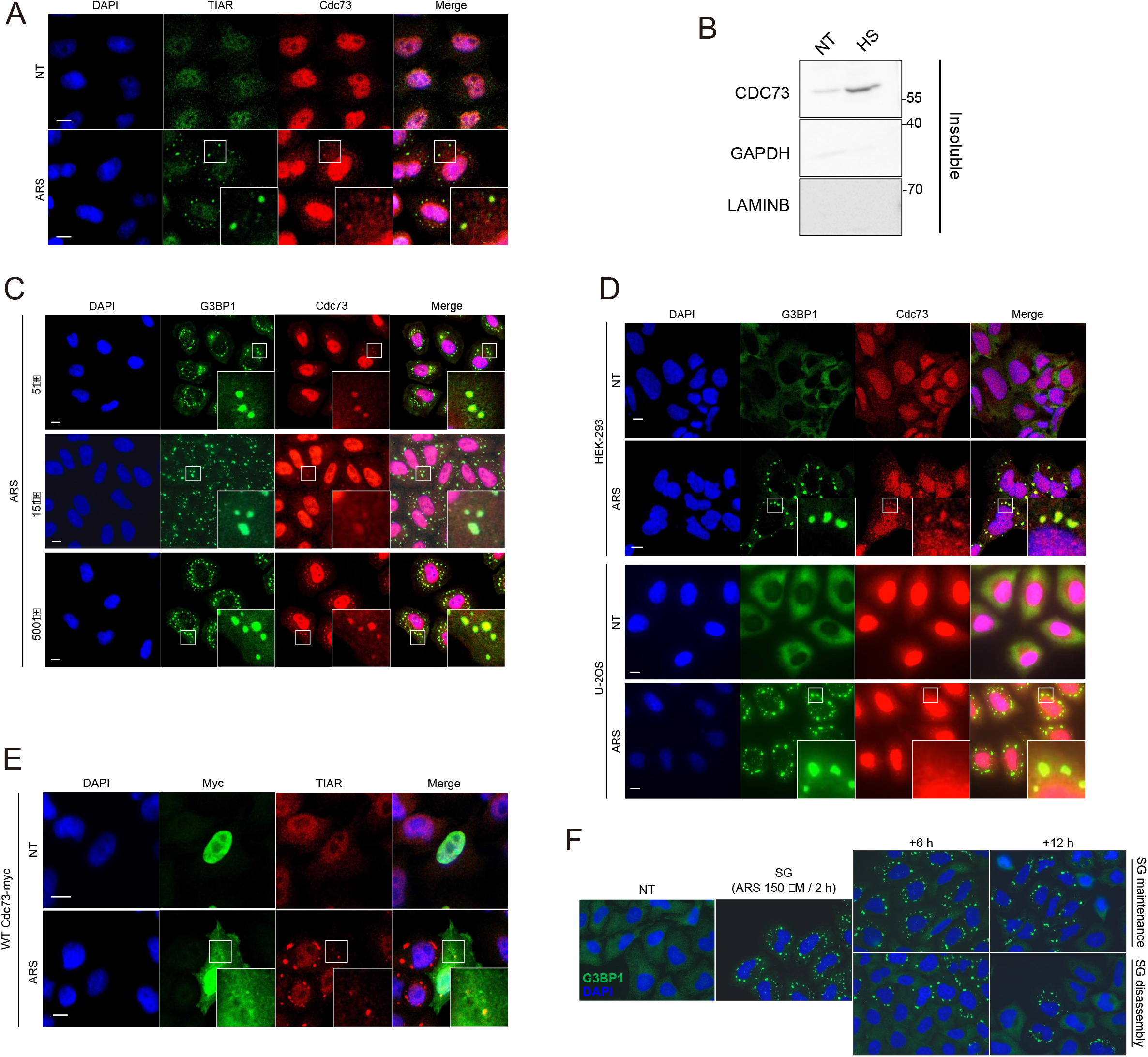
PDH LAMINB NT HS C Supplementary Figure S1. (A) Confocal images of endogenous hCdc73 and TIAR. HeLa cells were either untreated (NT) or treated with 500 μM sodium arsenite for 1 h (ARS). (B) Immunoblot analysis of hCdc73 from the insoluble fraction of cells exposed to heat shock (HS: 46°C for 1 h). (C) HeLa cells were treated with various concentrations of sodium arsenite (ARS: 50, 150, and 500 μM) for the indicated duration. Afterward, confocal images of endogenous hCdc73 and G3BP1 were taken. (D) Confocal images of HEK-293 and U-2OS cells treated with 500 μM sodium arsenite (ARS) for 1 h. (E) Confocal images of WT hCdc73-myc-transfected HeLa cells that were untreated (NT) or treated with 500 μM sodium arsenite for 1 h (ARS).(F) Representative immunofluorescence microscopy images of SG formation in HeLa cells. Endogenous G3BP1 and DAPI were stained after +6 h and +12 h of exposure to SG maintenance and disassembly conditions. The experimental scheme is shown in Figure 1G. G3BP1 and TIAR were used as markers for SGs. All scale bars in this figure, 10 μm. Each graph represents mean data from three independent experiments and error bars are presented as the mean ± standard deviations. Statistical values were calculated using an unpaired t-test.

**Supplementary Figure S2.**
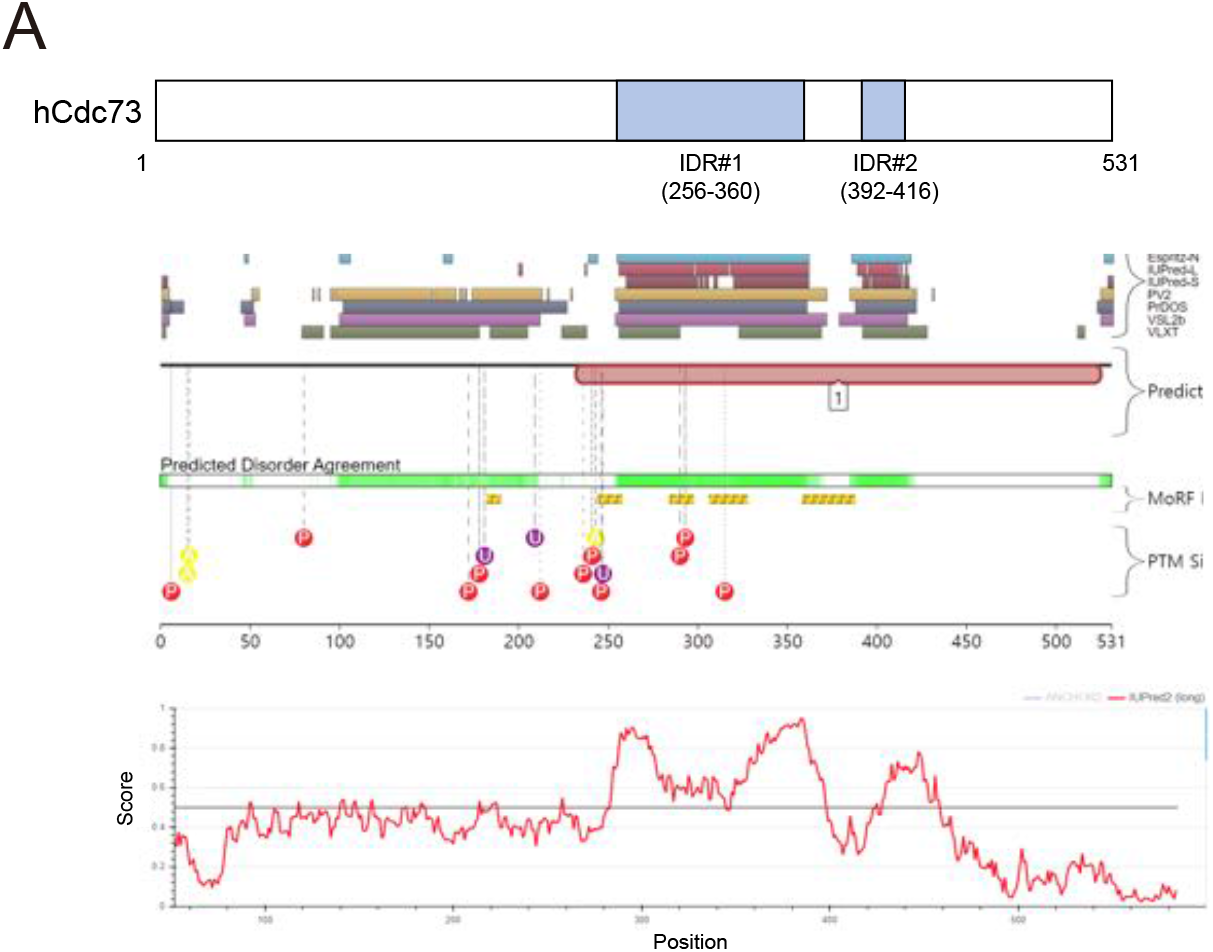
(A) (top) Schematic diagram of the mutation sites used to generate NLSmut hCdc73. (middle) Prediction of the IDR of hCdc73 from five different predictors in the D2P2 resource. (bottom) Presented IDR score of hCdc73 obtained from the IUPred simulation program. The threshold for all predictors was 0.5, and the disorder region is indicated as a gray box.

**Supplementary Figure S3.**
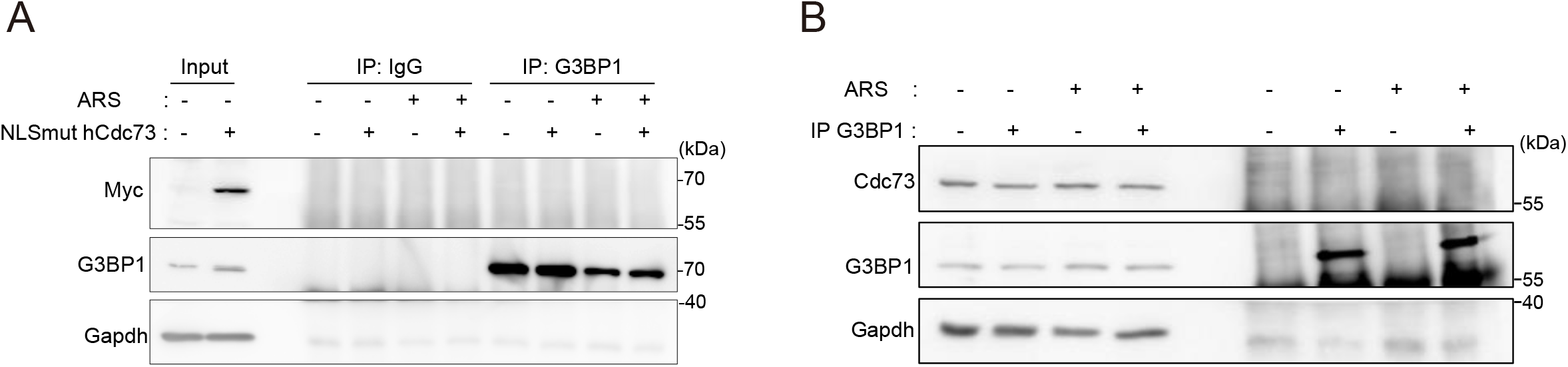
To cheak the interaction between G3BP1 and hCdc73, IP assay was performed with NLSmut hCdc73-myc (A) and endogenous hCdc73 (B) with G3BP1 in HeLa cells.

**Supplementary Figure S4.**
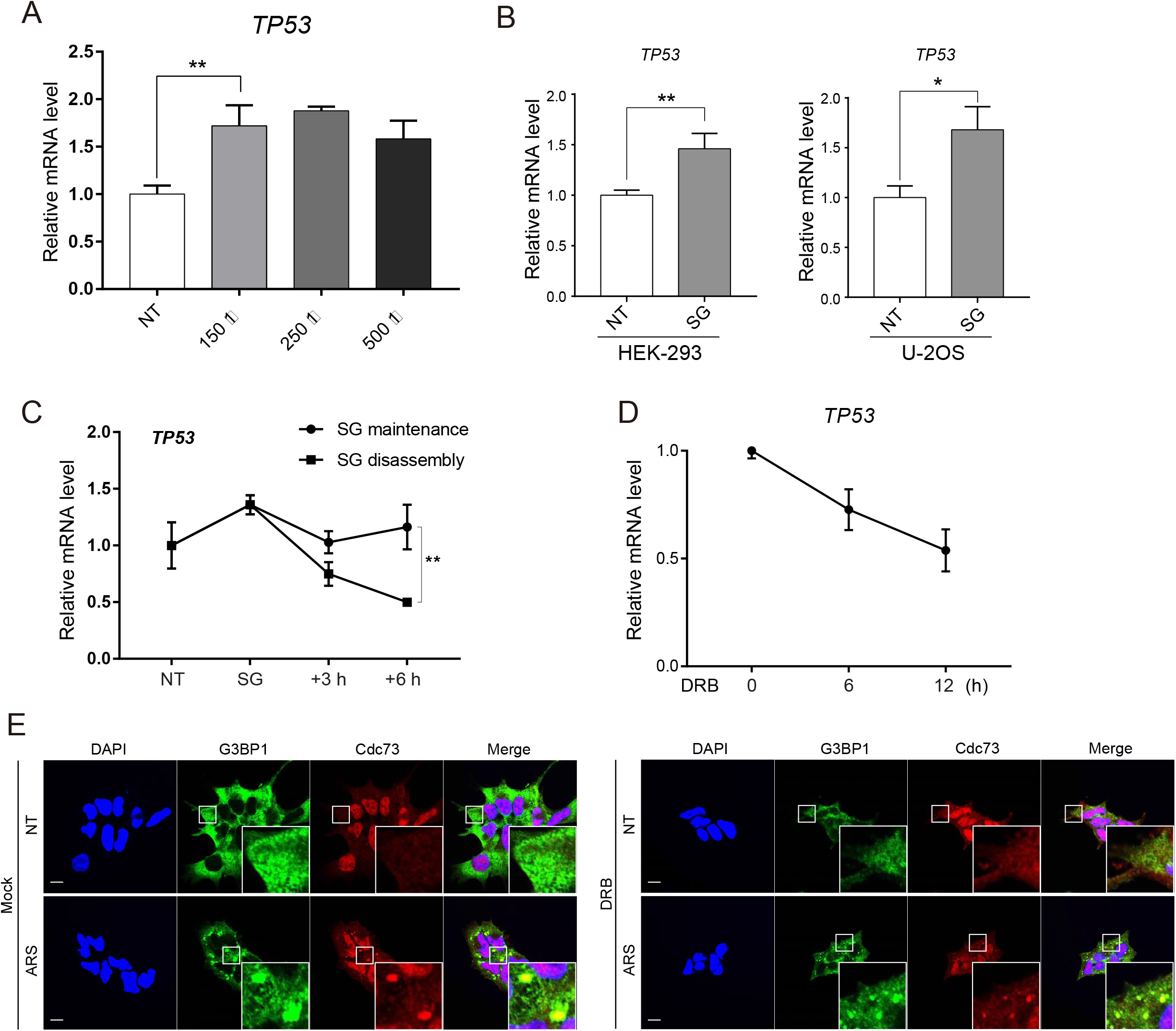
(A) HeLa cells were treated with 150, 250, or 500 μM sodium arsenite for 1 h. The p53 mRNA level in each stressed cell type was measured by qRT–PCR and compared with the NT (untreated) cells. (B) HEK-293 and U-2OS cells were stressed by treatment with 500 μM sodium arsenite for 1 h (SG). The p53 mRNA levels were measured by qRT–PCR. (C)HeLa cells were treated with 500 μM sodium arsenite for 1 h (SG) and then subjected to SG maintenance or disassembly according to the protocol shown in Figure 5B. At the indicated times, p53 mRNA levels were measured by qRT–PCR. (D) HEK-293 cells were treated with 100 μM DRB for 6 and 12 h. The p53 mRNA level was measured after qRT–PCR. (E) Confocal images of endogenous hCdc73 and G3BP1. HEK-293 cells were treated with DMSO (Mock) or 100 μM DRB for 12 h (DRB). After DRB treatment, 150 μM sodium arsenite was added, as shown in Figure 5C

**Supplementary Figure S5.**
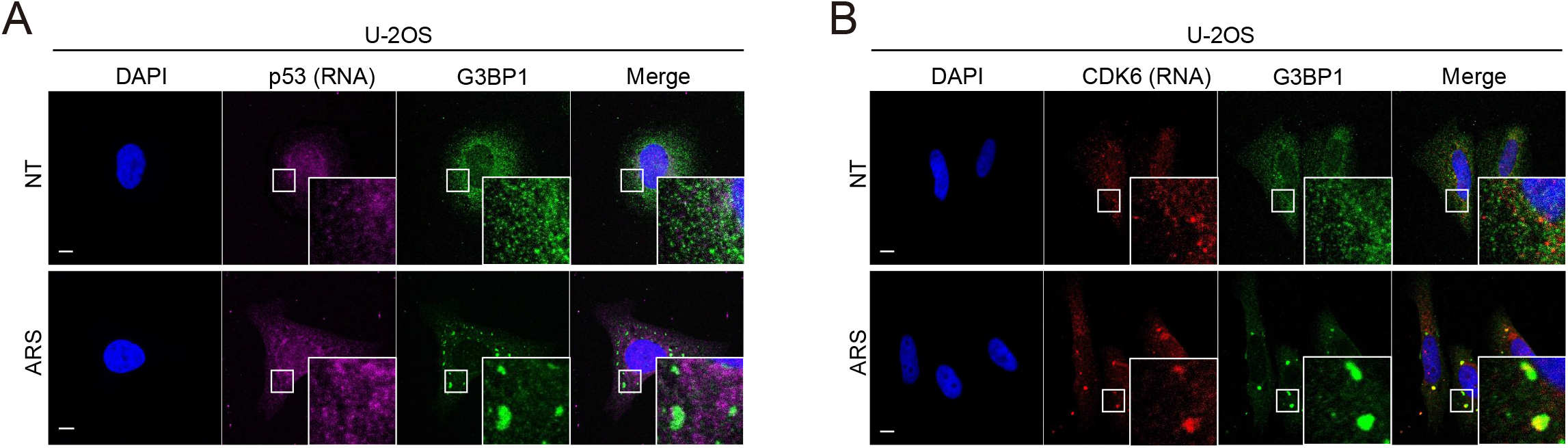
(A-B) Confocal images from RNA FISH experiments. U-2OS cells were either untreated (NT) or treated with 500 μM sodium arsenite for 1 h. FISH probes that specifically bind p53 (A) and Cdk6 (B) mRNAs were used. G3BP1 was used as a marker for SGs. Scale bars, 10 μm.

**Supplementary Table 1.**
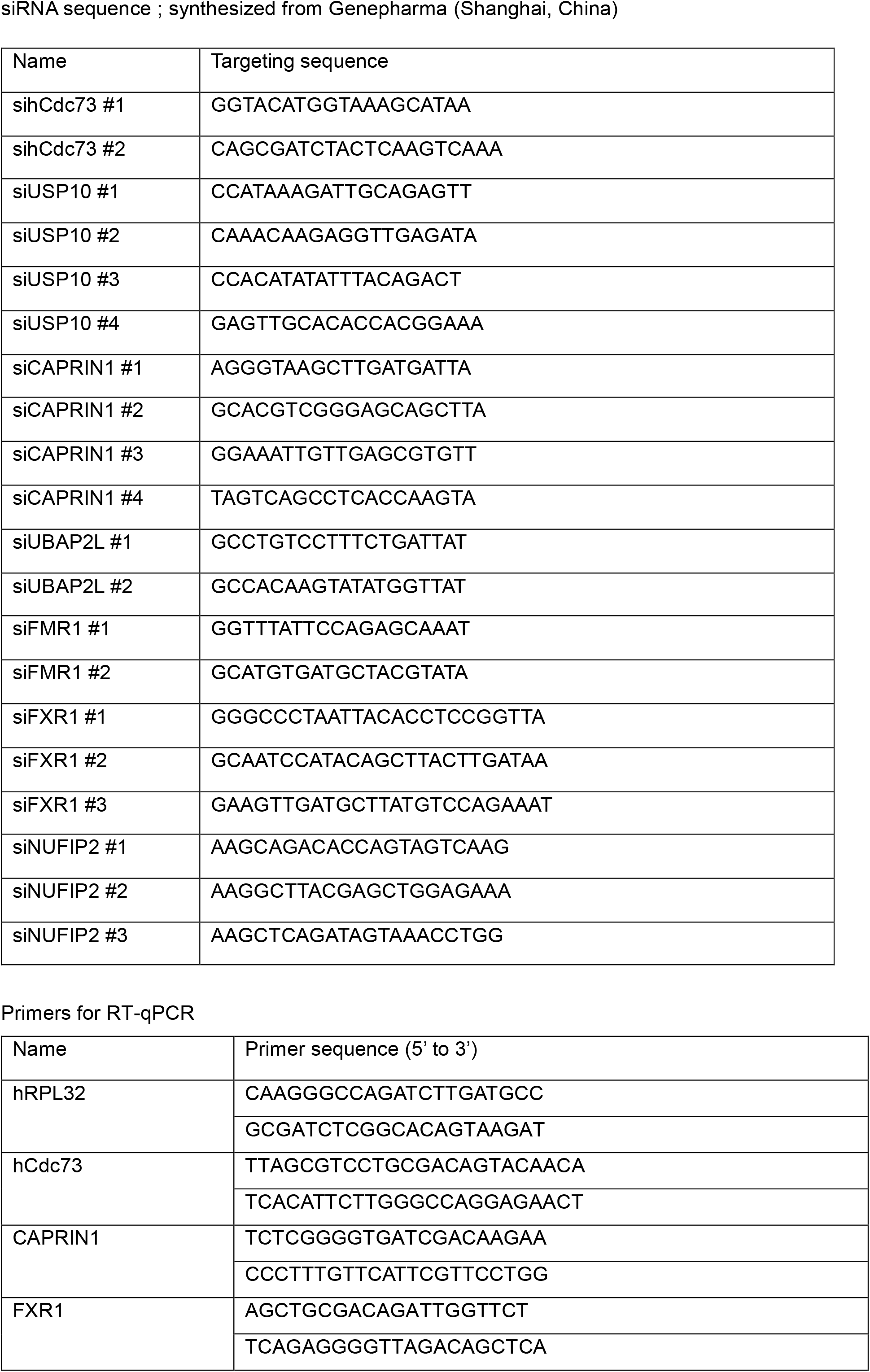
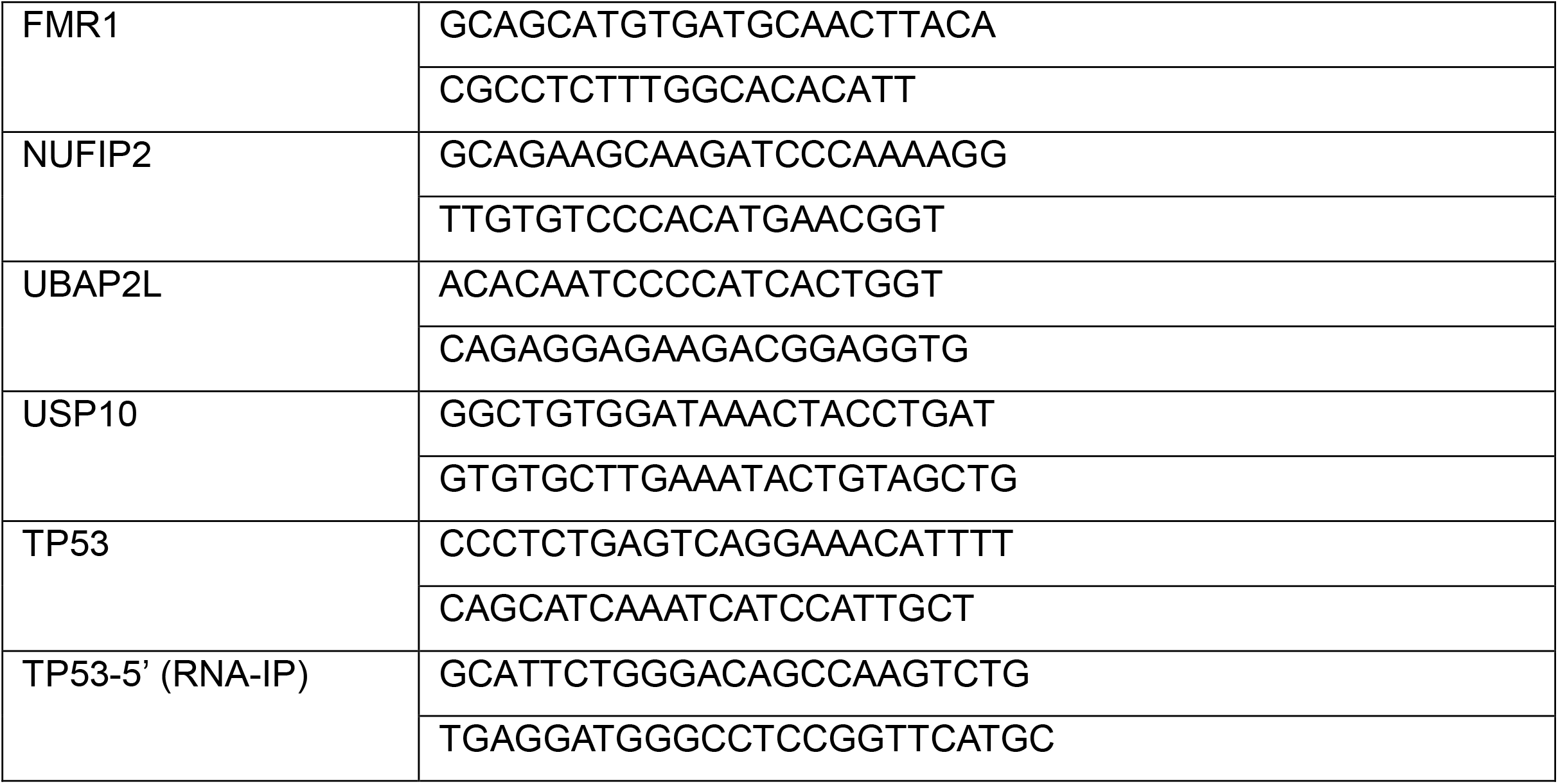
siRNA and primer sequence information used in this study

## REFERENCES

1. Kultz, D. (2005). Molecular and evolutionary basis of the cellular stress response. Annu Rev Physiol, 67, 225–257. https://doi.org/10.1146/annurev.physiol.67.040403.103635

2. Galluzzi, L., Yamazaki, T., & Kroemer, G. (2018). Linking cellular stress responses to systemic homeostasis. Nat Rev Mol Cell Biol, 19(11), 731–745. https://doi.org/10.1038/s41580-018-0068-0

3. Martindale, J. L., & Holbrook, N. J. (2002). Cellular response to oxidative stress: signaling for suicide and survival. J Cell Physiol, 192(1), 1–15. https://doi.org/10.1002/jcp.10119

4. Ciccia, A., & Elledge, S. J. (2010). The DNA damage response: making it safe to play with knives. Mol Cell, 40(2), 179–204. https://doi.org/10.1016/j.molcel.2010.09.019

5. Richter, K., Haslbeck, M., & Buchner, J. (2010). The heat shock response: life on the verge of death. Mol Cell, 40(2), 253–266. https://doi.org/10.1016/j.molcel.2010.10.006

6. Wang, Y. P., & Lei, Q. Y. (2018). Metabolite sensing and signaling in cell metabolism. Signal Transduct Target Ther, 3, 30. https://doi.org/10.1038/s41392-018-0024-7

7. Pizzinga, M., Harvey, R. F., Garland, G. D., Mordue, R., Dezi, V., Ramakrishna, M., Sfakianos, A., Monti, M., Mulroney, T. E., Poyry, T., & Willis, A. E. (2020). The cell stress response: extreme times call for post-transcriptional measures. Wiley Interdiscip Rev RNA, 11(3), e1578. https://doi.org/10.1002/wrna.1578

8. Fan, J., Yang, X., Wang, W., Wood, W. H., 3rd, Becker, K. G., & Gorospe, M. (2002). Global analysis of stress-regulated mRNA turnover by using cDNA arrays. Proc Natl Acad Sci U S A, 99(16), 10611–10616. https://doi.org/10.1073/pnas.162212399

9. Buchan, J. R. (2014). mRNP granules. Assembly, function, and connections with disease. RNA Biol, 11(8), 1019–1030. https://doi.org/10.4161/15476286.2014.972208

10. Shevtsov, S. P., & Dundr, M. (2011). Nucleation of nuclear bodies by RNA. Nat Cell Biol, 13(2), 167–173. https://doi.org/10.1038/ncb2157

11. Banani, S. F., Lee, H. O., Hyman, A. A., & Rosen, M. K. (2017). Biomolecular condensates: organizers of cellular biochemistry. Nat Rev Mol Cell Biol, 18(5), 285–298. https://doi.org/10.1038/nrm.2017.7

12. Protter, D. S. W., & Parker, R. (2016). Principles and Properties of Stress Granules. Trends Cell Biol, 26(9), 668–679. https://doi.org/10.1016/j.tcb.2016.05.004

13. Panas, M. D., Ivanov, P., & Anderson, P. (2016). Mechanistic insights into mammalian stress granule dynamics. J Cell Biol, 215(3), 313–323. https://doi.org/10.1083/jcb.201609081

14. Kimball, S. R., Horetsky, R. L., Ron, D., Jefferson, L. S., & Harding, H. P. (2003). Mammalian stress granules represent sites of accumulation of stalled translation initiation complexes. Am J Physiol Cell Physiol, 284(2), C273–284. https://doi.org/10.1152/ajpcell.00314.2002

15. Youn, J. Y., Dyakov, B. J. A., Zhang, J., Knight, J. D. R., Vernon, R. M., Forman-Kay, J. D., & Gingras, A. C. (2019). Properties of Stress Granule and P-Body Proteomes. Mol Cell, 76(2), 286–294. https://doi.org/10.1016/j.molcel.2019.09.014

16. Khong, A., Matheny, T., Jain, S., Mitchell, S. F., Wheeler, J. R., & Parker, R. (2017). The Stress Granule Transcriptome Reveals Principles of mRNA Accumulation in Stress Granules. Mol Cell, 68(4), 808–820 e805. https://doi.org/10.1016/j.molcel.2017.10.015

17. Van Treeck, B., Protter, D. S. W., Matheny, T., Khong, A., Link, C. D., & Parker, R. (2018). RNA self-assembly contributes to stress granule formation and defining the stress granule transcriptome. Proc Natl Acad Sci U S A, 115(11), 2734–2739. https://doi.org/10.1073/pnas.1800038115

18. Wilbertz, J. H., Voigt, F., Horvathova, I., Roth, G., Zhan, Y., & Chao, J. A. (2019). Single-Molecule Imaging of mRNA Localization and Regulation during the Integrated Stress Response. Mol Cell, 73(5), 946–958 e947. https://doi.org/10.1016/j.molcel.2018.12.006

19. Mateju, D., Eichenberger, B., Voigt, F., Eglinger, J., Roth, G., & Chao, J. A. (2020). Single-Molecule Imaging Reveals Translation of mRNAs Localized to Stress Granules. Cell, 183(7), 1801–1812 e1813. https://doi.org/10.1016/j.cell.2020.11.010

20. Jain, S., Wheeler, J. R., Walters, R. W., Agrawal, A., Barsic, A., & Parker, R. (2016). ATPase-Modulated Stress Granules Contain a Diverse Proteome and Substructure. Cell, 164(3), 487–498. https://doi.org/10.1016/j.cell.2015.12.038

21. Markmiller, S., Soltanieh, S., Server, K. L., Mak, R., Jin, W., Fang, M. Y., Luo, E. C., Krach, F., Yang, D., Sen, A., Fulzele, A., Wozniak, J. M., Gonzalez, D. J., Kankel, M. W., Gao, F. B., Bennett, E. J., Lecuyer, E., & Yeo, G. W. (2018). Context-Dependent and Disease-Specific Diversity in Protein Interactions within Stress Granules. Cell, 172(3), 590–604 e513. https://doi.org/10.1016/j.cell.2017.12.032

22. Youn, J. Y., Dunham, W. H., Hong, S. J., Knight, J. D. R., Bashkurov, M., Chen, G. I., Bagci, H., Rathod, B., MacLeod, G., Eng, S. W. M., Angers, S., Morris, Q., Fabian, M., Cote, J. F., & Gingras, A. C. (2018). High-Density Proximity Mapping Reveals the Subcellular Organization of mRNA-Associated Granules and Bodies. Mol Cell, 69(3), 517–532 e511. https://doi.org/10.1016/j.molcel.2017.12.020

23. Marmor-Kollet, H., Siany, A., Kedersha, N., Knafo, N., Rivkin, N., Danino, Y. M., Moens, T. G., Olender, T., Sheban, D., Cohen, N., Dadosh, T., Addadi, Y., Ravid, R., Eitan, C., Toth Cohen, B., Hofmann, S., Riggs, C. L., Advani, V. M., Higginbottom, A., … Hornstein, E. (2020). Spatiotemporal Proteomic Analysis of Stress Granule Disassembly Using APEX Reveals Regulation by SUMOylation and Links to ALS Pathogenesis. Mol Cell, 80(5), 876–891 e876. https://doi.org/10.1016/j.molcel.2020.10.032

24. Zhang, K., Daigle, J. G., Cunningham, K. M., Coyne, A. N., Ruan, K., Grima, J. C., Bowen, K. E., Wadhwa, H., Yang, P., Rigo, F., Taylor, J. P., Gitler, A. D., Rothstein, J. D., & Lloyd, T. E. (2018). Stress Granule Assembly Disrupts Nucleocytoplasmic Transport. Cell, 173(4), 958–971 e917. https://doi.org/10.1016/j.cell.2018.03.025

25. Vance, C., Scotter, E. L., Nishimura, A. L., Troakes, C., Mitchell, J. C., Kathe, C., Urwin, H., Manser, C., Miller, C. C., Hortobagyi, T., Dragunow, M., Rogelj, B., & Shaw, C. E. (2013). ALS mutant FUS disrupts nuclear localization and sequesters wild-type FUS within cytoplasmic stress granules. Hum Mol Genet, 22(13), 2676–2688. https://doi.org/10.1093/hmg/ddt117

26. Jaehning, J. A. (2010). The Paf1 complex: platform or player in RNA polymerase II transcription? Biochim Biophys Acta, 1799(5-6), 379–388. https://doi.org/10.1016/j.bbagrm.2010.01.001

27. Rozenblatt-Rosen, O., Hughes, C. M., Nannepaga, S. J., Shanmugam, K. S., Copeland, T. D., Guszczynski, T., Resau, J. H., & Meyerson, M. (2005). The parafibromin tumor suppressor protein is part of a human Paf1 complex. Mol Cell Biol, 25(2), 612–620. https://doi.org/10.1128/MCB.25.2.612-620.2005

28. Zhu, B., Mandal, S. S., Pham, A. D., Zheng, Y., Erdjument-Bromage, H., Batra, S. K., Tempst, P., & Reinberg, D. (2005). The human PAF complex coordinates transcription with events downstream of RNA synthesis. Genes Dev, 19(14), 1668–1673. https://doi.org/10.1101/gad.1292105

29. Van Oss, S. B., Cucinotta, C. E., & Arndt, K. M. (2017). Emerging Insights into the Roles of the Paf1 Complex in Gene Regulation. Trends Biochem Sci, 42(10), 788–798. https://doi.org/10.1016/j.tibs.2017.08.003

30. Vos, S. M., Farnung, L., Linden, A., Urlaub, H., & Cramer, P. (2020). Structure of complete Pol II-DSIF-PAF-SPT6 transcription complex reveals RTF1 allosteric activation. Nat Struct Mol Biol, 27(7), 668–677. https://doi.org/10.1038/s41594-020-0437-1

31. Takahashi, A., Tsutsumi, R., Kikuchi, I., Obuse, C., Saito, Y., Seidi, A., Karisch, R., Fernandez, M., Cho, T., Ohnishi, N., Rozenblatt-Rosen, O., Meyerson, M., Neel, B. G., & Hatakeyama, M. (2011). SHP2 tyrosine phosphatase converts parafibromin/Cdc73 from a tumor suppressor to an oncogenic driver. Mol Cell, 43(1), 45–56. https://doi.org/10.1016/j.molcel.2011.05.014

32. Kikuchi, I., Takahashi-Kanemitsu, A., Sakiyama, N., Tang, C., Tang, P. J., Noda, S., Nakao, K., Kassai, H., Sato, T., Aiba, A., & Hatakeyama, M. (2016). Dephosphorylated parafibromin is a transcriptional coactivator of the Wnt/Hedgehog/Notch pathways. Nat Commun, 7, 12887. https://doi.org/10.1038/ncomms12887

33. Rozenblatt-Rosen, O., Nagaike, T., Francis, J. M., Kaneko, S., Glatt, K. A., Hughes, C. M., LaFramboise, T., Manley, J. L., & Meyerson, M. (2009). The tumor suppressor Cdc73 functionally associates with CPSF and CstF 3’ mRNA processing factors. Proc Natl Acad Sci U S A, 106(3), 755–760. https://doi.org/10.1073/pnas.0812023106

34. Jo, J. H., Chung, T. M., Youn, H., & Yoo, J. Y. (2014). Cytoplasmic parafibromin/hCdc73 targets and destabilizes p53 mRNA to control p53-mediated apoptosis. Nat Commun, 5, 5433. https://doi.org/10.1038/ncomms6433

35. Pisani, C., Onori, A., Gabanella, F., Delle Monache, F., Borreca, A., Ammassari-Teule, M., Fanciulli, M., Di Certo, M. G., Passananti, C., & Corbi, N. (2016). eEF1Bgamma binds the Che-1 and TP53 gene promoters and their transcripts. J Exp Clin Cancer Res, 35(1), 146. https://doi.org/10.1186/s13046-016-0424-x

36. Marijan, D., Tse, R., Elliott, K., Chandhok, S., Luo, M., Lacroix, E., & Audas, T. E. (2019). Stress-specific aggregation of proteins in the amyloid bodies. FEBS Lett, 593(22), 3162–3172. https://doi.org/10.1002/1873-3468.13597

37. Boncella, A. E., Shattuck, J. E., Cascarina, S. M., Paul, K. R., Baer, M. H., Fomicheva, A., Lamb, A. K., & Ross, E. D. (2020). Composition-based prediction and rational manipulation of prion-like domain recruitment to stress granules. Proc Natl Acad Sci U S A, 117(11), 5826–5835. https://doi.org/10.1073/pnas.1912723117

38. Hahn, M. A., & Marsh, D. J. (2005). Identification of a functional bipartite nuclear localization signal in the tumor suppressor parafibromin. Oncogene, 24(41), 6241–6248. https://doi.org/10.1038/sj.onc.1208778

39. Hahn, M. A., & Marsh, D. J. (2007). Nucleolar localization of parafibromin is mediated by three nucleolar localization signals. FEBS Lett, 581(26), 5070–5074. https://doi.org/10.1016/j.febslet.2007.09.050

40. Amrich, C. G., Davis, C. P., Rogal, W. P., Shirra, M. K., Heroux, A., Gardner, R. G., Arndt, K. M., & VanDemark, A. P. (2012). Cdc73 subunit of Paf1 complex contains C-terminal Ras-like domain that promotes association of Paf1 complex with chromatin. J Biol Chem, 287(14), 10863–10875. https://doi.org/10.1074/jbc.M111.325647

41. Mosimann, C., Hausmann, G., & Basler, K. (2006). Parafibromin/Hyrax activates Wnt/Wg target gene transcription by direct association with beta-catenin/Armadillo. Cell, 125(2), 327–341. https://doi.org/10.1016/j.cell.2006.01.053

42. Newey, P. J., Bowl, M. R., Cranston, T., & Thakker, R. V. (2010). Cell division cycle protein 73 homolog (CDC73) mutations in the hyperparathyroidism-jaw tumor syndrome (HPT-JT) and parathyroid tumors. Hum Mutat, 31(3), 295–307. https://doi.org/10.1002/humu.21188

43. Sun, W., Kuang, X. L., Liu, Y. P., Tian, L. F., Yan, X. X., & Xu, W. (2017). Crystal structure of the N-terminal domain of human CDC73 and its implications for the hyperparathyroidism-jaw tumor (HPT-JT) syndrome. Sci Rep, 7(1), 15638. https://doi.org/10.1038/s41598-017-15715-9

44. Oates, M. E., Romero, P., Ishida, T., Ghalwash, M., Mizianty, M. J., Xue, B., Dosztanyi, Z., Uversky, V. N., Obradovic, Z., Kurgan, L., Dunker, A. K., & Gough, J. (2013). D(2)P(2): database of disordered protein predictions. Nucleic Acids Res, 41(Database issue), D508–516. https://doi.org/10.1093/nar/gks1226

45. Nott, T. J., Petsalaki, E., Farber, P., Jervis, D., Fussner, E., Plochowietz, A., Craggs, T. D., Bazett-Jones, D. P., Pawson, T., Forman-Kay, J. D., & Baldwin, A. J. (2015). Phase transition of a disordered nuage protein generates environmentally responsive membraneless organelles. Mol Cell, 57(5), 936–947. https://doi.org/10.1016/j.molcel.2015.01.013

46. Molliex, A., Temirov, J., Lee, J., Coughlin, M., Kanagaraj, A. P., Kim, H. J., Mittag, T., & Taylor, J. P. (2015). Phase separation by low complexity domains promotes stress granule assembly and drives pathological fibrillization. Cell, 163(1), 123–133. https://doi.org/10.1016/j.cell.2015.09.015

47. Shin, Y., Berry, J., Pannucci, N., Haataja, M. P., Toettcher, J. E., & Brangwynne, C. P. (2017). Spatiotemporal Control of Intracellular Phase Transitions Using Light-Activated optoDroplets. Cell, 168(1-2), 159–171 e114. https://doi.org/10.1016/j.cell.2016.11.054

48. Banani, S. F., Rice, A. M., Peeples, W. B., Lin, Y., Jain, S., Parker, R., & Rosen, M. K. (2016). Compositional Control of Phase-Separated Cellular Bodies. Cell, 166(3), 651–663. https://doi.org/10.1016/j.cell.2016.06.010

49. Sanders, D. W., Kedersha, N., Lee, D. S. W., Strom, A. R., Drake, V., Riback, J. A., Bracha, D., Eeftens, J. M., Iwanicki, A., Wang, A., Wei, M. T., Whitney, G., Lyons, S. M., Anderson, P., Jacobs, W. M., Ivanov, P., & Brangwynne, C. P. (2020). Competing Protein-RNA Interaction Networks Control Multiphase Intracellular Organization. Cell, 181(2), 306–324 e328. https://doi.org/10.1016/j.cell.2020.03.050

50. Taha, M. S., Haghighi, F., Stefanski, A., Nakhaei-Rad, S., Kazemein Jasemi, N. S., Al Kabbani, M. A., Gorg, B., Fujii, M., Lang, P. A., Haussinger, D., Piekorz, R. P., Stuhler, K., & Ahmadian, M. R. (2021). Novel FMRP interaction networks linked to cellular stress. FEBS J, 288(3), 837–860. https://doi.org/10.1111/febs.15443

51. Riback, J. A., Katanski, C. D., Kear-Scott, J. L., Pilipenko, E. V., Rojek, A. E., Sosnick, T. R., & Drummond, D. A. (2017). Stress-Triggered Phase Separation Is an Adaptive, Evolutionarily Tuned Response. Cell, 168(6), 1028–1040 e1019. https://doi.org/10.1016/j.cell.2017.02.027

52. Kruiswijk, F., Labuschagne, C. F., & Vousden, K. H. (2015). p53 in survival, death and metabolic health: a lifeguard with a licence to kill. Nat Rev Mol Cell Biol, 16(7), 393–405. https://doi.org/10.1038/nrm4007

53. Kruse, J. P., & Gu, W. (2009). Modes of p53 regulation. Cell, 137(4), 609–622. https://doi.org/10.1016/j.cell.2009.04.050

54. Diaz-Munoz, M. D., Kiselev, V. Y., Le Novere, N., Curk, T., Ule, J., & Turner, M. (2017). Tia1 dependent regulation of mRNA subcellular location and translation controls p53 expression in B cells. Nat Commun, 8(1), 530. https://doi.org/10.1038/s41467-017-00454-2

55. Takayama, K. I., Suzuki, T., Fujimura, T., Takahashi, S., & Inoue, S. (2018). Association of USP10 with G3BP2 Inhibits p53 Signaling and Contributes to Poor Outcome in Prostate Cancer. Mol Cancer Res, 16(5), 846–856. https://doi.org/10.1158/1541-7786.MCR-17-0471

56. Boija, A., Klein, I. A., Sabari, B. R., Dall’Agnese, A., Coffey, E. L., Zamudio, A. V., Li, C. H., Shrinivas, K., Manteiga, J. C., Hannett, N. M., Abraham, B. J., Afeyan, L. K., Guo, Y. E., Rimel, J. K., Fant, C. B., Schuijers, J., Lee, T. I., Taatjes, D. J., & Young, R. A. (2018). Transcription Factors Activate Genes through the Phase-Separation Capacity of Their Activation Domains. Cell, 175(7), 1842–1855 e1816. https://doi.org/10.1016/j.cell.2018.10.042

57. Shrinivas, K., Sabari, B. R., Coffey, E. L., Klein, I. A., Boija, A., Zamudio, A. V., Schuijers, J., Hannett, N. M., Sharp, P. A., Young, R. A., & Chakraborty, A. K. (2019). Enhancer Features that Drive Formation of Transcriptional Condensates. Mol Cell, 75(3), 549–561 e547. https://doi.org/10.1016/j.molcel.2019.07.009

58. Lu, Y., Wu, T., Gutman, O., Lu, H., Zhou, Q., Henis, Y. I., & Luo, K. (2020). Phase separation of TAZ compartmentalizes the transcription machinery to promote gene expression. Nat Cell Biol, 22(4), 453–464. https://doi.org/10.1038/s41556-020-0485-0

59. Lu, H., Yu, D., Hansen, A. S., Ganguly, S., Liu, R., Heckert, A., Darzacq, X., & Zhou, Q. (2018). Phase-separation mechanism for C-terminal hyperphosphorylation of RNA polymerase II. Nature, 558(7709), 318–323. https://doi.org/10.1038/s41586-018-0174-3

60. Dosztányi, Z., Csizmók, V., Tompa, P., & Simon, I. (2005). The Pairwise Energy Content Esti mated from Amino Acid Composition Discriminates between Folded and Intrinsically Unstruct ured Proteins. Journal of Molecular Biology, 347(4), 827–839. https://doi.org/10.1016/j.jmb.2005.01.071

